# The lack of macrophage fragment adhesion is a benchmark of dormant hematopoietic stem cells throughout the lifespan

**DOI:** 10.64898/2026.06.29.735148

**Authors:** Masashi Kanayama, Yuta Izumi, Yuki Yamada, Satoko Arakawa, Atsushi Iwama, Toshiaki Ohteki

**Affiliations:** Department of Biodefense Research, Medical Research Laboratory, Institute of Integrated Research, Institute of Science Tokyo (formerly Medical Research Institute, Tokyo Medical and Dental University (TMDU)), Tokyo, Japan; Ochanomizu Research Facility (ORF), Bioscience Center, Research Infrastructure Management Center, Institute of Science Tokyo (formerly Medical Research Institute, Tokyo Medical and Dental University (TMDU)), Tokyo, Japan; Division of Stem Cell and Molecular Medicine, Center for Stem Cell Biology and Regenerative Medicine, The Institute of Medical Science, The University of Tokyo, Tokyo, Japan

## Abstract

Hematopoietic stem cells (HSCs) play a pivotal role in the lifelong maintenance of hematopoiesis. However, heterogeneity and age-related alterations in HSC populations hinders accurate HSC analysis. Here, we show that bone marrow (BM) macrophage fragments that preferentially express F4/80 adhere to proliferative rather than dormant HSCs. The adhesion of macrophage fragments to proliferative HSCs occurred throughout the process of BM cell preparation *in vitro*. Consistently, proliferative HSCs express genes involved in the adhesion of macrophage fragments at higher levels than dormant HSCs. Notably, by using that as a benchmark, dormant HSCs can be easily identified as F4/80^low^HSCs throughout their lifespan, thereby revealing that they retain considerable stemness and remain functional with aging. Collectively, we propose a novel and straightforward method for the rapid identification, isolation, and analysis of distinct HSC subpopulations, which will be helpful for a wide range of hematological studies and will provide insights into HSC biology. (149 words)

## Introduction

Hematopoietic stem cells (HSCs) play a crucial role in maintaining homeostasis and influencing the pathogenesis of multiple diseases by producing blood cells, including red blood cells, platelets, and immune cells. Therefore, elucidating how HSCs are maintained and altered during different life stages and disease progression is essential to understand the evolution of life and the pathogenesis of diseases. However, the HSC fraction is heterogeneous, and biological stresses can alter the population size and functionality of dormant HSCs, a subpopulation of HSCs with high stemness^1, 2^. Due to the inaccuracy of identifying HSC subpopulations, it has been challenging to determine whether the results reflect intrinsic alterations in HSCs or compositional changes within HSC subpopulations. To address that issue, a side-population identification method was previously developed based on the drug efflux characteristic of dormant HSCs^3^. Detecting nitric oxide or cytoplasmic calcium also identified dormant HSCs^4, 5^. However, those methods require the culture of HSCs, which carries the risk of altering their functions, gene expression patterns, and/or stage of differentiation. Therefore, the development of a method to quickly identify HSC subpopulations without the requirement for their culture has been sought. Single-cell RNA sequencing enables the precise detection of dormant HSCs, but the RNA extraction process kills those cells, making it difficult to isolate the target cells. To easily identify and isolate dormant HSCs, their specific surface markers have been proposed^5, 6, 7, 8, 9, 10^. However, their utility under biological stresses has not been fully elucidated.

A variety of cells and molecules are involved in the construction of the HSC niche^11, 12^. Especially, CXCR4, a receptor for CXCL12, plays significant roles in quantitatively and qualitatively maintaining HSCs by receiving CXCL12 produced by cells that constitute the HSC niche^13, 14, 15^. It has recently been reported that HSCs acquire CXCR4 from bone marrow (BM) macrophages through trogocytosis, a phenomenon in which membrane molecules of one cell are functionally acquired through endocytosis by another cell, thereby retaining themselves in the BM^16^. On the other hand, another report suggested that fragments of BM macrophages adhered to hematopoietic stem progenitor cells (HSPCs), including HSCs, during cell preparation^17^. However, the former study did not rigorously demonstrate trogocytosis by HSCs, and the latter study did not confirm the physiological significance of macrophage fragment adhesion, leading to potential confusion in HSC biology.

This study demonstrates that BM macrophage fragments preferentially attach to proliferative, rather than dormant, HSCs *in vitro*. The attachment of macrophage fragments was easily detected by the surface expression of macrophage-associated molecules, including F4/80 and CD169. While the expression of F4/80 was not detected on the cell surface of HSCs when they were recruited into the blood, where BM macrophages do not exist, F4/80 became detectable when HSCs were mixed with a BM cell suspension that contains BM macrophages *in vitro*. Additionally, the macrophage fragments were not functionally integrated with the cell membrane of HSCs. Notably, the absence of macrophage fragment attachments was a consistent feature of dormant HSCs throughout their lifespan, thereby revealing that they retain considerable stemness and remain functional with aging. Taken together, this new method to quickly identify dormant HSCs using macrophage fragment adhesion as an indicator minimizes the confounding effect of HSC heterogeneity and enables the accurate analysis of HSC subpopulations even under stress conditions.

## Results

### Adhesion of macrophage fragments to hematopoietic progenitors

In mixed BM chimeras generated by transferring BM cells obtained from *CAG-EGFP* mice and tamoxifen-treated *Rosa26-lsl-tdTomato; Rosa26-CreERT2* mice (Fig. 1a), some EGFP^high^ and tdTomato^high^ cells in the long-term HSCs (LT-HSCs) fraction (Supplementary Fig. 1a) expressed intermediate levels of tdTomato and EGFP, respectively (Fig. 1b). Confocal microscopic analysis revealed that those intermediate expression levels reflected the attachment of EGFP-expressing cell fragments on the surface of tdTomato^high^ LT-HSCs (Fig. 1c). In this context, using electron microscopy (EM), we observed 170 cell fragments on HSPCs and found that all of them were attached to the cell surface. In addition, no clathrin structure formation or endocytic uptake was observed at the interface with the cell fragments (Fig. 1d and Supplementary Fig. 1b-d).

**Fig. 1:**
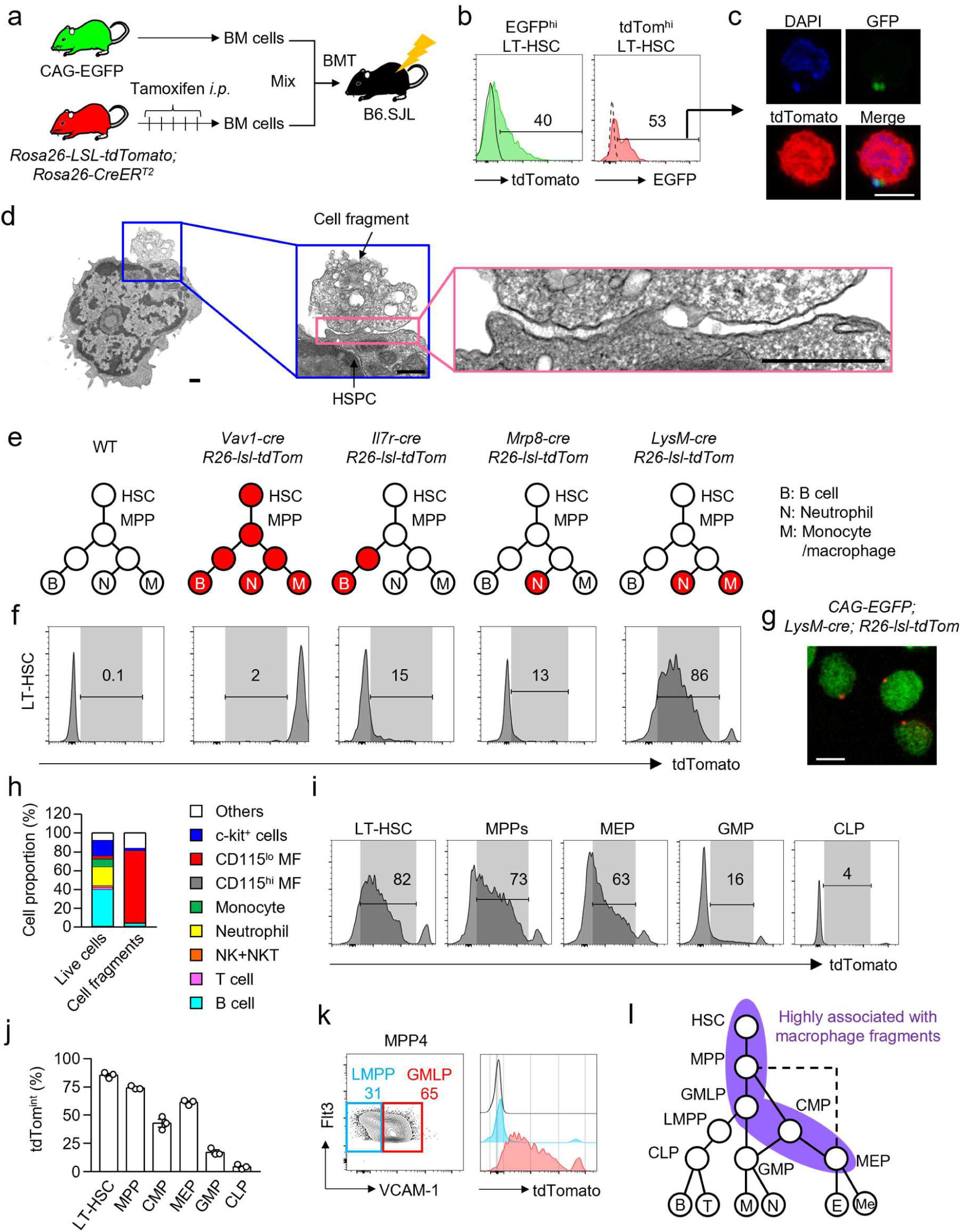
Macrophage fragments adhere to the surface of HSCs. **a-c** Detection of macrophage fragments on the surface of LT-HSCs. Tamoxifen (2 mg/mouse/day) was intraperitoneally injected into *Rosa26-lsl-tdTomato; Rosa26-CreERT2* mice for five consecutive days. One to four weeks after the last tamoxifen treatment, BM cells obtained from the mice were mixed with the BM cells of CAG-EGFP mice and were transferred into lethally irradiated SJL mice. Four weeks after the transplantation, the expression levels of tdTomato in EGFP^high^ LT-HSCs and of EGFP in tdTomato^high^ LT-HSCs were examined by FACS (b). EGFP^+^ tdTomato^high^ LT-HSCs were sorted and analyzed by confocal microscopy (c). The experimental strategy is shown in (a). **d** EM analysis of cell fragments attached to HSPCs. Scale bars indicate 0.5 μm. **e, f** Expression of tdTomato in LT-HSCs obtained from WT, *Vav1-Cre; Rosa26-lsl-tdTomato*, *Il7r-Cre; Rosa26-lsl-tdTomato*, *Mrp8-Cre; Rosa26-lsl-tdTomato* and *LysM-Cre; Rosa26-lsl-tdTomato* mice. Schematic illustration of tdTomato-expressing cells (red circles) is shown in (e), and representative FACS plots are shown in (f). Numbers in the FACS plots indicate tdTomato^int^ cell frequencies (shown as gray areas in f). **g** Confocal microscopy analysis of tdTomato^int^ LT-HSCs obtained from *CAG-EGFP; LysM-Cre; Rosa26-lsl-tdTomato* mice. **h** Cell proportion in live cell population and cell fragments in the BM of naïve WT mice. **i-l** Expression of tdTomato^int^ cells (shown as gray areas in FACS plots) in LT-HSCs, MPPs, MEPs, GMPs, CLPs (i, j) and LMPPs and GMLPs (k) in the BM of *LysM-Cre; Rosa26-lsl-tdTomato* mice. In (k), as a control, tdTomato expression in WT MPP4s is shown (unfilled histogram). The purple area in the schematic illustration of the hematopoietic differentiation pathway indicates cells showing higher frequencies of tdTomato^int^ cells in *LysM-Cre; Rosa26-lsl-tdTomato* mice in (l). B: B cell, T: T cell, M: monocyte/macrophage, N: neutrophil, E: erythrocyte, Me: megakaryocyte. Data represent two (c, d, g-k) or three (b, f, i) independent experiments.

To examine what type of cell fragments attach to LT-HSCs, we prepared *Vav1-cre; Rosa26-lsl-tdTomato* mice (labeling all blood cells), *Il7r-cre; Rosa26-lsl-tdTomato* mice (labeling lymphoid cells), *Mrp8-cre; Rosa26-lsl-tdTomato* mice (labeling neutrophils), and *LysM-cre; Rosa26-lsl-tdTomato* mice (labeling myeloid cells) (Fig. 1e). LT-HSCs intrinsically expressing tdTomato were detected as tdTomato^high^ in *Vav1-cre; Rosa26-lsl-tdTomato* mice, and those with cell fragment adhesion were detected as intermediate expression of tdTomato (tdTomato^int^, indicated as gray areas) (Fig. 1f). We found that tdTomato^int^ LT-HSCs were present in *LysM-cre; Rosa26-lsl-tdTomato* mice, but not in *Il7r-cre; Rosa26-lsl-tdTomato* mice or *Mrp8-cre; Rosa26-lsl-tdTomato* mice (Fig. 1f, g), suggesting that the cell fragments were derived from myeloid cells other than neutrophils. In this context, most cell fragments (CD45^+^FSC^low^) were derived from CD115^low^ macrophages (Fig. 1h and Supplementary Fig. 1e, f), also known as CD169^+^ BM-resident macrophages^18^. Notably, macrophages were a minor fraction both in live and in dead cells (Fig. 1h and Supplementary Fig. 1g), which suggests that BM CD115^low^ macrophages are prone to fragmentation. Conversely, what type of hematopoietic progenitors do the fragments bind to? In addition to LT-HSCs, macrophage fragments were attached to multipotent progenitors (MPPs), common myeloid progenitors (CMPs), and megakaryocyte-erythrocyte progenitors (MEPs), but not granulocyte-macrophage progenitors (GMPs) or common lymphoid progenitors (CLPs) (Fig. 1i, j). Among the MPP4 fraction, macrophage fragments were attached to granulocyte/monocyte/lymphoid progenitors (GMLPs) but not to lympho-myeloid primed progenitors (LMPPs) (Fig. 1k). Thus, the macrophage fragments preferentially adhered to the cell surface of multipotent and erythroid progenitors (Fig. 1l).

### Detection of macrophage markers on HSCs

In the mixed BM chimeras generated using BM cells obtained from SJL (CD45.1^+^CD45.2^-^) mice and CAG-EGFP mice (CD45.1^-^CD45.2^+^) (Supplementary Fig. 2a), CD45.1 expression was detected on the surface of LT-HSCs from CAG-EGFP mice (Supplementary Fig. 2b). Those expression levels were correlated with the chimerism of SJL-derived cells (Supplementary Fig. 2c), suggesting that CD45.1 was acquired from SJL mouse-derived cells. We analyzed the expression of 253 molecules on the surface of HSPCs of WT mice (the gating strategy is shown in Supplementary Fig. 1a). Notably, the expression levels of 24 molecules including macrophage-related markers were significantly correlated with F4/80, a well-established marker of macrophages, suggesting that those molecules were possibly derived from macrophage fragments (Supplementary Fig. 2d). While the macrophage-related markers F4/80, CD169, CD64, CD14, and VCAM-1, were clearly detected on the surface of LT-HSCs (Supplementary Fig. 2e), mRNA expression levels of the genes encoding those molecules (*Adgre1*, *Siglec1*, *Fcgr1*, *Cd14*, and *Vcam1*) were as weak as those of negative control genes such as *Ighm* encoding immunoglobulin, *Cryab* encoding lens structural protein, and *Krt32* encoding Keratin (Supplementary Fig. 2e, f). These results suggested that the acquisition of macrophage markers occurred at the protein level rather than at the mRNA level. Consistently, the cell surface levels of F4/80 and CD169 on LT-HSCs were correlated with the amounts of macrophage fragments attached to the LT-HSCs (Supplementary Fig. 2g). In addition, the levels of cell fragments on LMPPs and GMLPs correlated well with the expression levels of F4/80 (Fig. 1j and Supplementary Fig. 2h). In this context, the expression levels of *Adgre1*, which encodes F4/80, were low both in LMPPs and in GMLPs, suggesting that the cell surface expression of F4/80 on GMLPs reflected the attachment of macrophage fragments (Supplementary Fig. 2i). Collectively, the attachment of macrophage fragments can be easily identified by detecting the expression of macrophage markers such as F4/80 and CD169 on HSPCs and HSCs.

Next, we investigated whether macrophage fragments adhere to HSPCs *in vivo*. First, we mixed CD169 knock out (KO) HSPCs with a BM cell suspension from WT mice on ice and confirmed the expression of CD169 on the surface of those CD169 KO HSPCs (Fig. 2a-c). Thus, the *in vitro* mixing was sufficient to induce the adhesion of macrophage fragments to HSPCs. It has been reported that HSPCs that emerged in the blood after treatment with G-CSF or AMD3100, a CXCR4 inhibitor, did not express cell surface F4/80^16^. We hypothesized that the failure to detect F4/80^+^ HSPCs could be simply due to the absence of BM macrophage fragments in the blood (Supplementary Fig. 2j). To examine that possibility, we induced the egress of HSPCs by administering G-CSF or AMD3100 into WT mice (Fig. 2d). As expected, no blood HSPCs expressed F4/80 (Fig. 2e, f). However, when the blood F4/80^-^HSPCs were mixed with the WT BM cell suspension on ice, the majority of them gained the expression of F4/80 (Fig. 2e, f). Conversely, treating HSPCs and LT-HSCs with trypsin-EDTA significantly decreased the frequencies of F4/80^high^ cells. At the same time, the expression levels of Sca-1 and c-kit were unaltered (Fig. 2g, h), suggesting that most macrophage fragments were not fused with HSPCs. Supporting this finding, a previous study using imaging flow cytometry observed F4/80 expression localized to attached macrophage fragments but not to the plasma membrane of HSCs and MPPs^17^. Consistently, immune electron microscopy (IEM) analysis demonstrated that F4/80 was expressed in the cell fragments but not on the cell membrane of HSPCs (Fig. 2i and Supplementary Fig. 2k-n). Based on these findings, we concluded that the macrophage fragment adhesion predominantly occurs *in vitro* (Supplementary Fig. 2j).

**Fig. 2:**
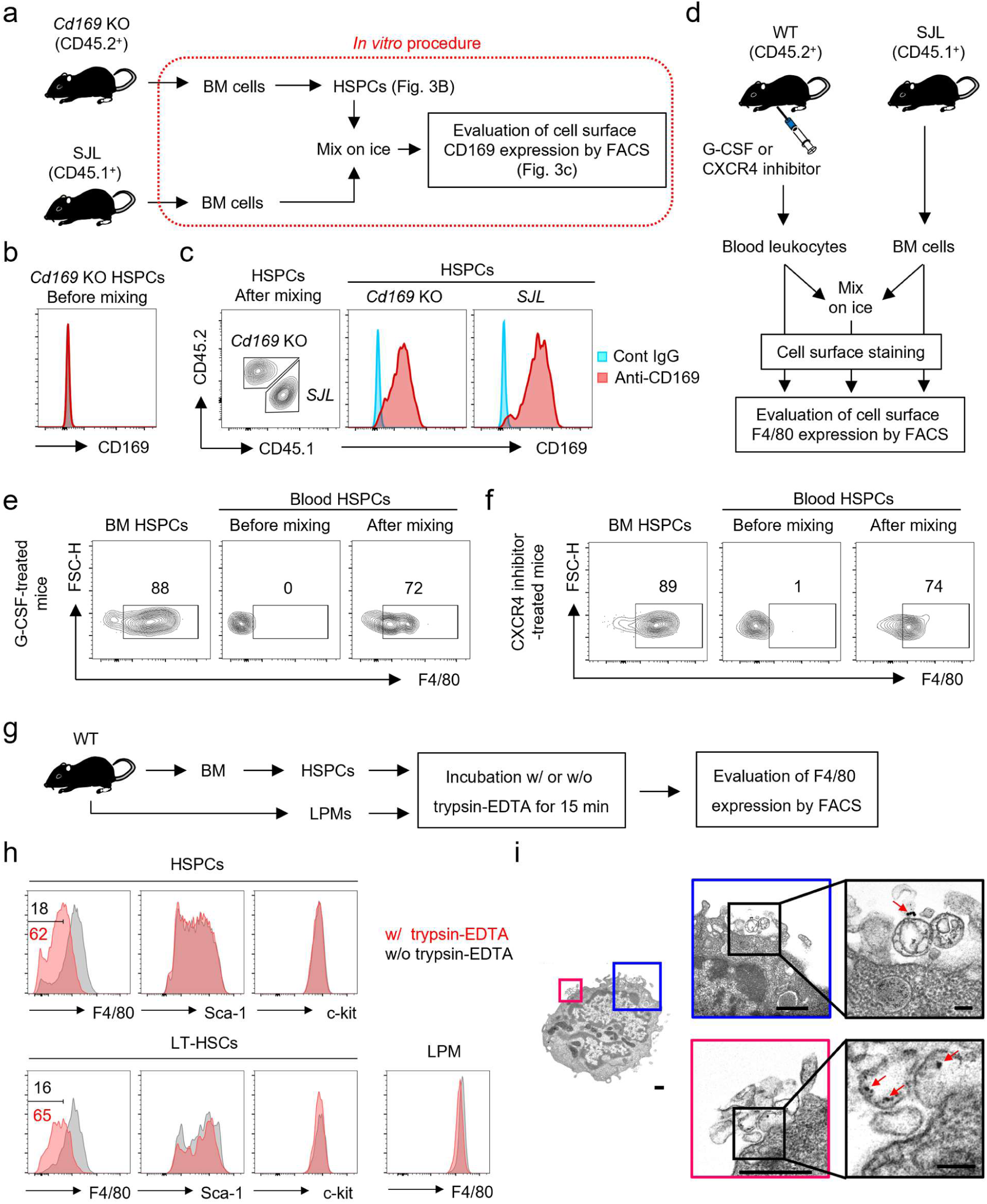
Macrophage fragments adhere to HSPCs *in vitro*. **a-c** HSPCs obtained from *Cd169*-deficient mice (CD45.2^+^) were mixed with a BM cell suspension obtained from SJL mice (CD45.1^+^) on ice and the expression of CD169 on the surface of *Cd169*-deficient HSPCs was evaluated before (b) and after the mixing (c). The experimental strategy is shown in (a). **d-f** Blood cells obtained from WT mice (CD45.2^+^) treated with G-CSF (e) or a CXCR4 inhibitor (f) were mixed with a BM cell suspension from SJL mice (CD45.1^+^). The expression of F4/80 was evaluated in the blood HSPCs before and after the mixing. The experimental strategy is shown in (d). **g, h** Treatment with trypsin-EDTA reduced F4/80 expression in HSPCs. HSPCs and large peritoneal macrophages (LPM) obtained from naïve WT B6 mice were treated with 0.03% trypsin-EDTA at 37°C for 15 min after which the expression of F4/80 was examined by FACS. The experimental strategy is shown in (g). **i** Immune electron microscopy of F4/80 detected on the surface of HSPCs obtained from the BM of naïve WT mice. Red arrows indicate F4/80 expression detected by gold particles. Scale bars in the rightmost panels indicate 0.1 μm and scale bars in the other panels indicate 0.5 μm. Data are representative of two independent experiments (a-c, e, f, h).

### Macrophage-derived molecules on HSCs are dysfunctional

The CXCL12/CXCR4 axis is the primary mechanism by which HSCs home into the BM and it supports the dormancy of HSCs within the BM niche. In this context, a recent report suggested that HSCs acquire CXCR4 from BM macrophages through trogocytosis, a process in which membrane molecules of one cell are functionally acquired by another cell, thereby retaining themselves in the BM^16^. To examine that, we tested the functions of macrophage-associated molecules on HSPCs and on HSCs. While the expression levels of cell surface CXCR4 protein on F4/80^high^ HSPCs were much higher than those on F4/80^lo^ HSPCs (Supplementary Fig. 2d), mRNA expression levels of *Cxcr4* in F4/80^high^ HSPCs were significantly lower than in F4/80^low^ HSPCs (Fig. 3a), which implies that the majority of cell surface CXCR4 on F4/80^high^ HSPCs is macrophage fragment-derived CXCR4. Indeed, the migration ability of F4/80^high^ HSPCs toward CXCL12 was significantly lower than that of F4/80^low^ HSPCs (Fig. 3b), reflecting the expression level of intrinsic CXCR4. These results suggested that the macrophage fragment-derived CXCR4 was not functional.

**Fig. 3:**
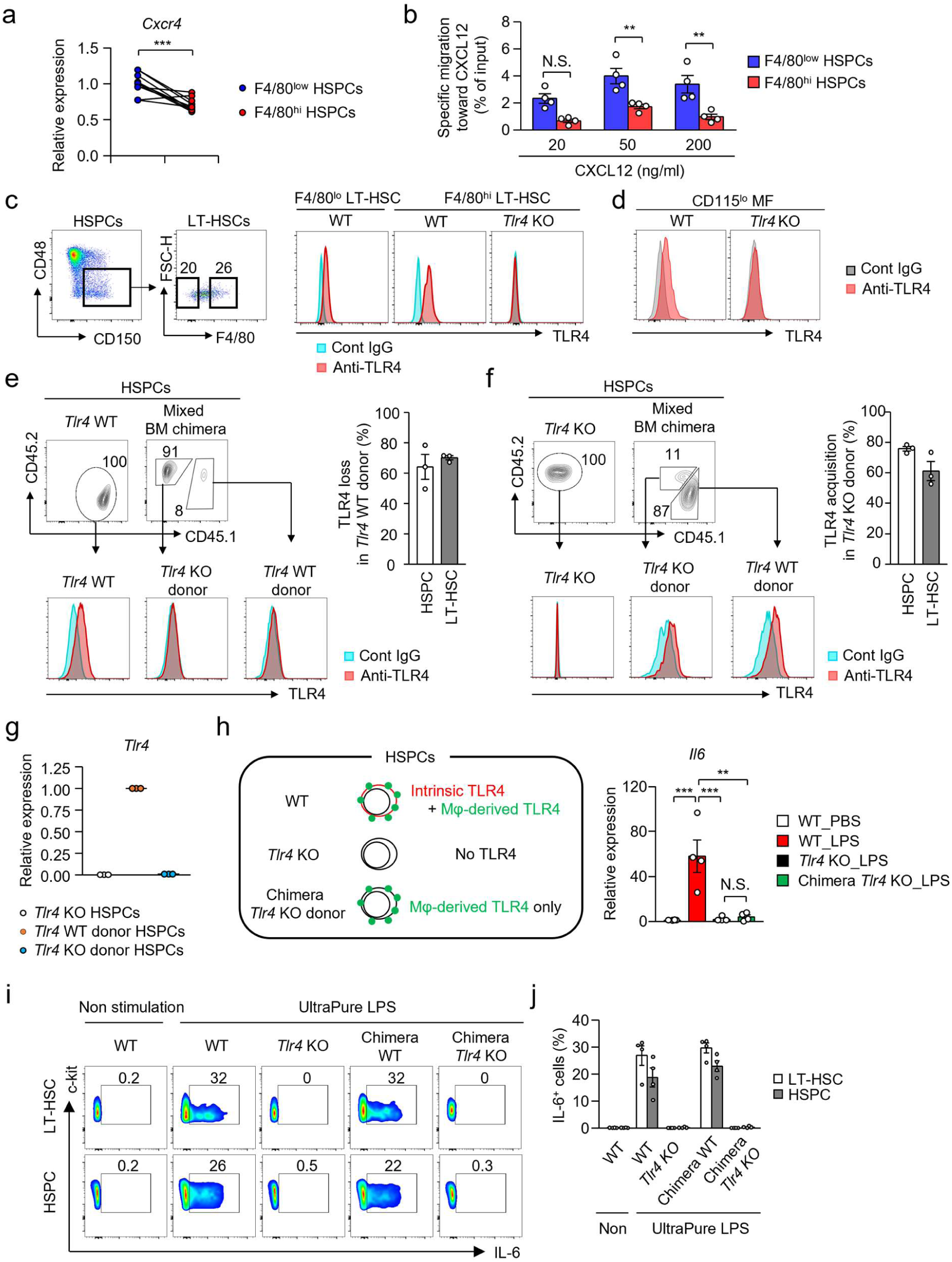
Macrophage-derived cell surface molecules do not signal to HSCs. **a** The expression level of *Cxcr4* mRNA in F4/80^low^ and in F4/80^high^ HSPCs was evaluated using qPCR. n=10 per group. **b** Transwell migration assays of F4/80^low^ and F4/80^high^ HSPCs to CXCL12. F4/80^low^ and F4/80^high^ HSPCs were obtained from the BM of naïve WT mice and were added in the upper chamber, and recombinant mouse CXCL12 was added to the bottom chamber at the indicated concentration. Migrated cell numbers were counted after 24 hours of culture. n=4 per group. **c** Expression of TLR4 on the surface of F4/80^low^ and of F4/80^high^ LT-HSCs. **d** Expression of TLR4 on CD115^low^ BM macrophages. **e** BM cells of *Tlr4* WT (SJL, CD45.1^+^) and *Tlr4* cKO (*Tlr4* ^flox/flox^; *Vav1*-cre, CD45.2^+^) mice were mixed at a 1:9 ratio and transferred into recipient *Tlr4* cKO mice as shown in Supplementary Fig. 4a. The loss of TLR4 in SJL donors was calculated by comparing TLR4 mean fluorescence intensity (MFI) in SJL donors between before and after transplantation (right). n=3 per group. **f** BM cells of *Tlr4* WT (CD45.1^+^) mice and *Tlr4* cKO (CD45.2^+^) mice were mixed at a 9:1 ratio and transferred into recipient SJL mice, as shown in Supplementary Fig. 4b. The acquisition of TLR4 in the *Tlr4* cKO donor was calculated by comparing TLR4 MFI between the *Tlr4* cKO donor and the *Tlr4* WT donor (right). n=3 per group. **g** *Tlr4* WT HSPCs and cKO donor HSPCs were obtained from BM chimera mice as shown in Supplementary Fig. 3b and the expression of *Tlr4* was examined by qPCR. **h** *Tlr4* cKO donor HSPCs were isolated as shown in Supplementary Fig. 3b and were stimulated with ultrapure LPS (100 μg/ml) for 6 hours after which the upregulation of *Il6* was evaluated. n=4 per group. **i, j** *Tlr4* WT HSPCs and cKO donor HSPCs were obtained from BM chimera mice as shown in Supplementary Fig. 3b and were stimulated with ultrapure LPS (100 μg/ml) for 6 hours after which the production of IL-6 in HSPCs and LT-HSCs was examined. Error bars, SEM. **p<0.01, *** p<0.001, N.S.; not significant. Student’s t-test [a], one-way ANOVA was used [b, h]. Data are representative of two (c, d) or four (i) independent experiments or were pooled from two (a), three (e-g) or four (b, j) independent experiments.

To further clarify whether HSCs functionally acquire cell surface molecules from macrophage fragments, we focused on TLR4, which is known to activate HSCs^19^. Like other macrophage-derived molecules, TLR4 was expressed at higher levels on F4/80^high^ LT-HSCs than on F4/80^low^ LT-HSCs (Fig. 3c), implying that some TLR4 molecules on the surface of F4/80^high^ LT-HSCs reflect the attachment of macrophage fragments derived from CD115^low^ BM macrophages that expressed TLR4 (Fig. 3d). To definitively prove that, we generated mixed BM chimeras by injecting *Tlr4^flox/flox^; Vav1-Cre* (hereafter, referred to as *Tlr4* KO) and *Tlr4* WT BM cells at a 9:1 ratio (Supplementary Fig. 3a), and found that *Tlr4* WT LT-HSCs and HSPCs lost about 70% of cell surface TLR4 expression (Fig. 3e). On the contrary, in mixed BM chimeras generated by transplanting *Tlr4* cKO and *Tlr4* WT BM cells at a 1:9 ratio (Supplementary Fig. 3b), *Tlr4* KO LT-HSCs and HSPCs acquired approximately 70% of the TLR4 expression level of WT LT-HSCs and HSPCs (Fig. 3f). Considering the selective adhesion of macrophage fragments (Fig. 1h and Supplementary Fig. 1e), these results indicated that the majority of cell surface TLR4 molecules on HSPCs and LT-HSCs of WT mice were derived from BM macrophages. In this context, *Tlr4* KO HSPCs that acquired cell-surface TLR4 did not express *Tlr4* mRNA (Fig. 3g). Lastly, we stimulated the TLR4-bearing *Tlr4* KO HSPCs with ultrapure lipopolysaccharide (LPS) (Supplementary Fig. 3b). While WT HSPCs, which express both intrinsic and macrophage-derived TLR4, upregulated their expression of *Il6* mRNA upon LPS stimulation, *Tlr4* KO HSPCs, which express only macrophage-derived TLR4, did not (Fig. 3h). Similar results were obtained from experiments with TLR4-bearing *Tlr4* KO LT-HSCs (Fig. 3i, j). Taken together, these results indicate that the macrophage fragments on HSPCs and HSCs are dysfunctional, which further suggests that trogocytosis is unlikely to be involved in this process.

### Phenotypical and functional characteristics of F4/80^high^ and F4/80^low^ HSCs

Based on the amounts of macrophage fragments adhering to HSCs, HSCs could be classified into F4/80^high^ HSCs and F4/80^low^ HSCs. Thus, we explored the significance of this phenomenon. In CD34^-^HSPCs, where HSCs are enriched, the frequency of CD48^-^CD150^+^ LT-HSCs^20^ was significantly enriched in the F4/80^low^ fraction compared to the F4/80^high^ fraction (Fig. 4a, B). In this context, we realized that F4/80^low^ LT-HSCs had a lower expression of CD48 and c-kit, and a higher expression of Sca-1 (Fig. 4a, c, d), which is a characteristic of HSCs with a higher BM reconstitution capacity^21, 22^. Consistently, F4/80^low^ LT-HSCs contained far more CD34^-^ cells than F4/80^high^ LT-HSCs (Fig. 4e, f), suggesting the abundance of dormant HSCs in the F4/80^low^ LT-HSC fraction.

**Fig. 4:**
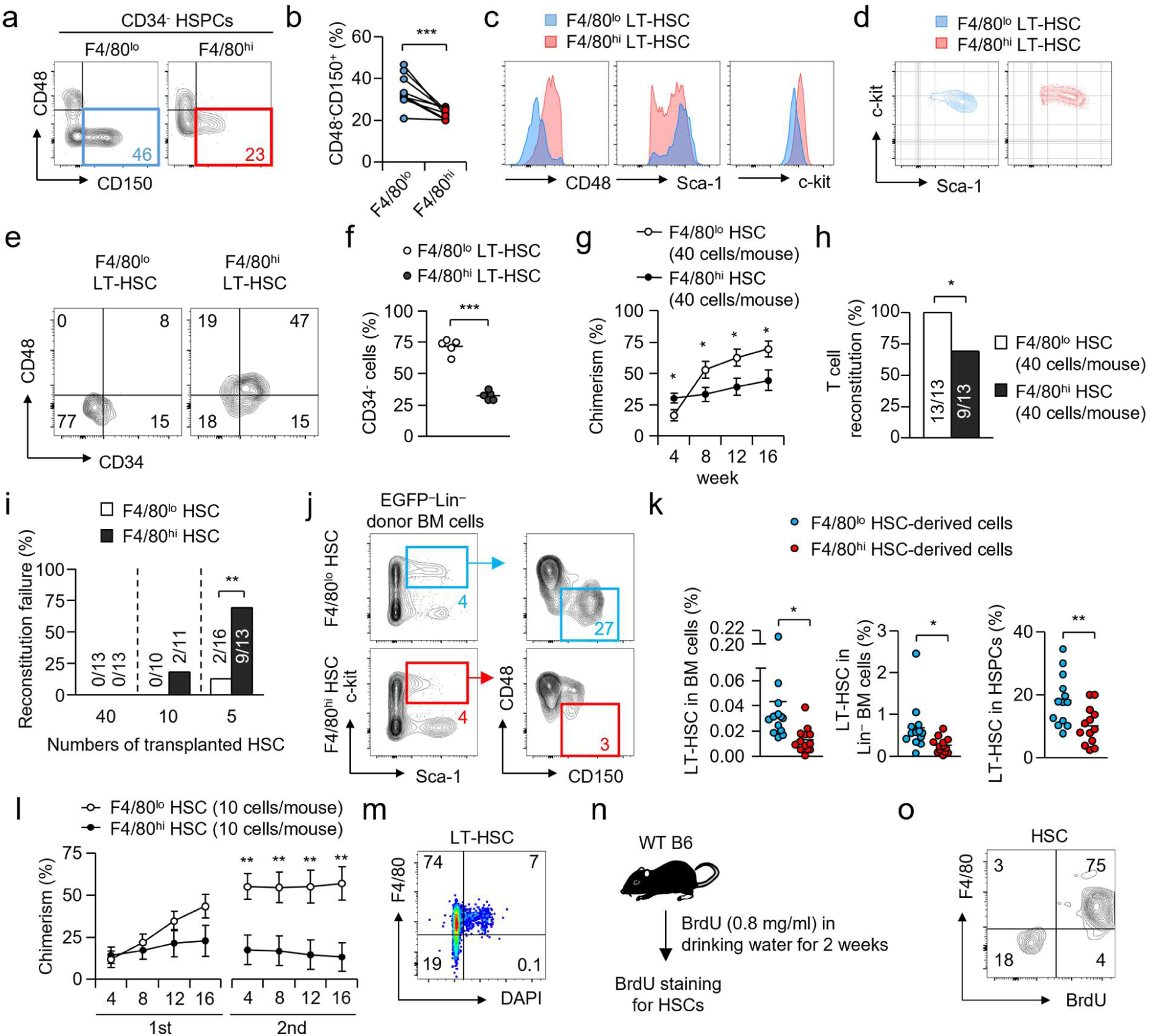
F4/80^low^ HSCs show higher BM reconstitution capacity and dormancy. **a, b** Frequency of CD48^-^CD150^+^ fraction in F4/80^low^ and F4/80^high^ fractions of CD34^-^HSPCs. Statistical evaluation is shown in (b). n=9 per group. **c, d** Expression of CD48, Sca-1 and c-kit in F4/80^low^ and in F4/80^high^ LT-HSCs (CD48^-^CD150^+^HSPCs). n=5 per group. **e, f** Expression of CD34 in F4/80^low^ and in F4/80^high^ LT-HSCs. Statistical evaluation is shown in (f). n=5 per group. **g-k** F4/80^low^ or F4/80^high^ HSCs were transplanted into lethally irradiated SJL mice with EGFP^+^ rescue BM cells (1 x 10^5^), as shown in Supplementary Fig. 4a. For (g, h, j, k), HSCs (40 cells) were transplanted (n=13 per group). Chimerism of F4/80^low^ or F4/80^high^ HSCs-derived cells in the blood is shown in (g), and the percentage of chimera mice that successfully reconstituted the T cell fraction is shown in (h). The frequencies of reconstitution failure at 16 weeks are shown in (i). The population size of F4/80^low^ or F4/80^high^ HSCs-derived LT-HSCs is shown in (j, k). **l** Serial transplantation of F4/80^low^ or F4/80^high^ HSCs. Ten F4/80^low^ or F4/80^high^ HSCs were transplanted into lethally irradiated SJL mice with EGFP^+^ rescue BM cells (1 x 10^5^). Sixteen weeks later, 2 x 10^7^ BM cells obtained from the recipient mice were transferred into another irradiated SJL mouse. The chimerism of F4/80^low^ or F4/80^high^ HSC-derived cells in blood leukocytes was examined every 4 weeks. n=10 for F4/80^low^ HSCs, n=11 for F4/80^high^ HSCs. **m** Detection of proliferating cells in LT-HSCs of naïve WT mice by DAPI staining. **n, o** BrdU incorporation by HSCs. WT mice were treated with BrdU for 2 weeks, and the incorporation of BrdU in HSCs was examined by FACS analysis. The experimental strategy is shown in (o). Error bars, SEM. *p<0.05, **p<0.01, ***p<0.001. Student’s t-test [b, f, g, k, l] and χ^2^ statistics [h, i] were used. Data are representative of two (e, m, n), three (c-h, j), or nine (a) independent experiments or were pooled from three (f-i, k, l) or nine (b) independent experiments.

To compare the BM reconstitution capacity of F4/80^low^ and F4/80^high^ CD34^-^LT-HSCs (hereafter CD34^-^LT-HSCs are referred to as HSCs, the gating strategy is shown in Supplementary Fig. 1a). We transferred F4/80^low^ or F4/80^high^ HSCs, along with EGFP-expressing rescue BM cells, into lethally irradiated SJL mice (Supplementary Fig. 4a). While the chimerism of F4/80^low^ HSC-derived cells was significantly lower than that of F4/80^high^ HSC-derived cells at 4 weeks after the transplantation (Fig. 4g), the chimerism of F4/80^low^ HSC-derived cells exceeded that of F4/80^high^ HSC-derived cells after 8 weeks. At 16 weeks, F4/80^low^ HSCs could reconstitute multiple lineages including T cells, B cells, neutrophils, and monocytes (Supplementary Fig. 4b) in all recipient mice, whereas F4/80^high^ HSCs failed to reconstitute the T cell lineage in about 30% of recipient mice (Fig. 4h and Supplementary Fig. 4b). Compared with F4/80^high^ HSCs, F4/80^low^ HSCs exhibited a lower rate of reconstitution failure, even with small numbers of HSCs (Fig. 4i). Notably, F4/80^low^ HSCs had a higher capacity to reconstitute the HSC pool compared to F4/80^high^ HSCs (Fig. 4j, k), resulting in more profound chimerism in the secondary transplantation than in the primary transplantation (Fig. 4l).

We further tested the dormancy of F4/80^low^ HSCs. DAPI staining of LT-HSCs revealed that the F4/80^high^ fraction, but not the F4/80^low^ fraction, contained cells in the G2/M phase (Fig. 4m). WT mice were treated with BrdU for 2 weeks, after which the incorporation status of BrdU in HSCs was examined. Most F4/80^high^ HSCs became BrdU^+^, whereas BrdU^+^ cells were hardly seen in F4/80^low^ HSCs (Fig. 4n, o). Collectively, these results strongly suggest that F4/80^high^ cells are proliferative HSCs, while F4/80^low^ cells are dormant HSCs.

### Single cell multiome analysis of F4/80^low^ and F4/80^high^ HSCs

To further support our findings, we performed CITE-seq analysis using LT-HSCs (Lin^-^Sca-1^+^c-kit^+^CD86^+^CD48^-^CD150^+^) from naïve WT mice. LT-HSCs were classified into 5 clusters based on antibody-capture profiles (Fig. 5a, b). Consistent with FACS profile and RNA-seq data (Supplementary Fig. 2d-f), the cell surface expression of macrophage markers such as F4/80, CD169, CD115, Clec12a, CD106, and MERTK was detected (Fig. 5c and Supplementary Fig. 5). In contrast, those molecules were not detected at the mRNA level (Fig. 5c and Supplementary Fig. 5). Compared with other clusters, cluster 4 exhibited a higher expression of Sca-1 and a lower expression of c-kit, CD34, and CD48 at the surface protein level (Fig. 5d), and lower expression of *Cd34* and an enrichment of *Procr*, which encodes EPCR, a marker for HSCs^23^, at the mRNA level (Fig. 5e). Notably, the cells in cluster 4 were relatively enriched with *Egr1*, *Ifitm1*, *Ltb*, *Cdkn1c*, *Junb*, *Fos*, *Cd74*, *Slfn2*, *Meg3*, and *Igf2bp2*, genes that promote dormancy or inhibit proliferation of HSCs^24, 25, 26, 27, 28, 29, 30, 31, 32, 33^ (shown in blue in Fig. 5e, f) and were relatively reduced with *Myc, Top2a*, *Spc24,* and *Mki67*, genes promoting cell cycle progression (shown in red in Fig. 5e, g). In line with our findings, *Myc* is known to be down-regulated in dormant HSCs^34, 35^. Consistent with the prominent multipotency of F4/80^low^ HSCs (Fig. 4k), cluster 4 weakly expressed cell surface CD41, a marker for myeloid and megakaryocyte-biased HSCs^36, 37^ (Fig. 5h). Since CD41^-^ HSCs were mostly dormant in naïve conditions^8^, this result also supported the dormancy of F4/80^low^ HSCs. In cluster 4, the expression of *Gfi1*, a transcription factor that promotes HSC self-renewal^38^ was relatively high (Fig. 5i), whereas the expression of *Phf6*, *Tet2*, *Dnmt3a,* and *Spread1*, genes that prevent self-renewal^38, 39, 40^, was relatively low (Fig. 5j). Notably, the expression of these gene sets in F4/80^high^ HSCs was opposite to that in F4/80^low^ HSCs. Thus, CITE-seq data strongly supported the classification of F4/80^low^ and F4/80^high^ HSCs as dormant and proliferative HSCs, respectively.

**Fig. 5:**
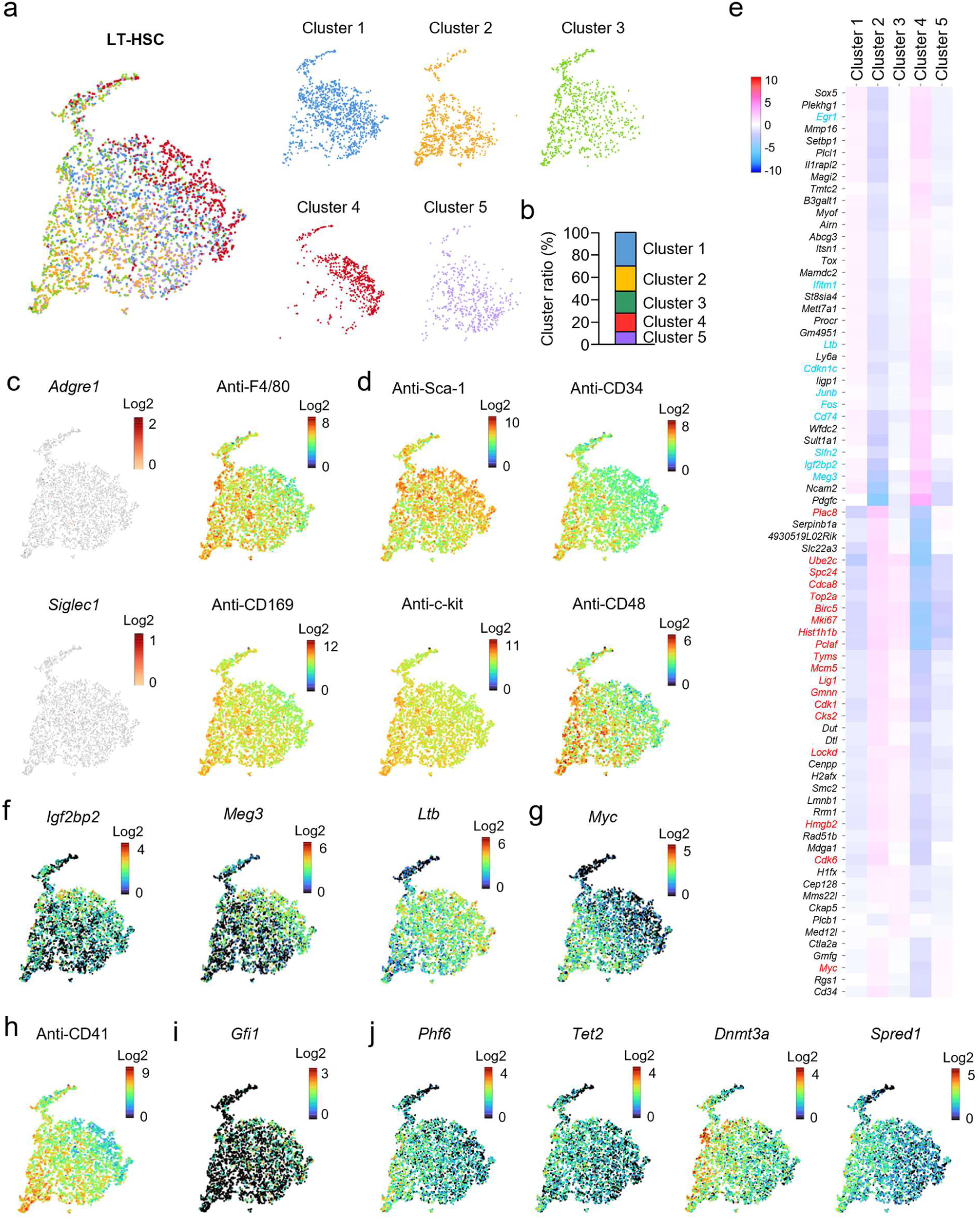
Enrichment of dormancy-related genes in F4/80^low^ HSCs. **a, b** Clustering of LT-HSCs (Lin^-^Sca-1^+^c-kit^+^CD86^+^CD48^-^CD150^+^ BM cells) obtained from naïve WT mice based on the expression profiles of cell surface molecules. The ratio of each cluster is shown in (b). **c** Expression of F4/80 and CD169 at the mRNA level (left) and at the cell surface protein level (right). **d** Expression of Sca-1 and c-kit and CD34 and CD48 on the surface of LT-HSCs. **e** Heatmap for differential expression genes of each cluster. Genes related to dormancy of HSCs are shown in blue and cell cycle-related genes are shown in red. **f, g** Expression of dormancy-related genes (f) and *Myc* (g). **h** Expression of CD41 on the surface of LT-HSCs. **i, j** Expression of genes that promote self-renewal (i) and genes that prevent self-renewal (j) in LT-HSCs.

### Mechanisms of macrophage fragment adhesion to proliferative HSCs

By what mechanism do macrophage fragments attach to proliferative HSCs? EM analysis implied the presence of specific molecules mediating the attachment of macrophage fragments to HSPCs (pink arrows in Supplementary Fig. 6a, b). Since a CD169-deficiency decreases the frequency of F4/80-expressing HSPCs^17^, we firstly examined whether CD169 directly mediates the attachment of macrophage fragments to LT-HSCs. We harvested BM cells in the presence or absence of a blocking antibody for CD169 and found that CD169 blockade partially but significantly decreased the frequency of the F4/80^+^ fraction (Supplementary Fig. 6c-e). Consistently, the expression level of *Spn*, which encodes CD43, a primary ligand for CD169^41^, was low in dormant HSCs (cluster 4) (Supplementary Fig. 6f). These results suggested that macrophage fragments attach to HSCs, at least in part, through the binding of CD169 on macrophage fragments to CD43 on F4/80^high^ HSCs during BM cell preparation. Second, the *in vitro* attachment of macrophage fragments to HSCs is prevented by cytochalasin D^16^, an inhibitor of actin polymerization. Notably, dormant HSCs (cluster 4) exhibited the lowest expression of *Actb*, the gene encoding β-actin (Fig. 6a). In addition, the expression of genes that are critically involved in actin rearrangements, such as *Rac1*, *Rac2*, *Cdc42*, *Rhoa*, and *Pik3ca*, integrins, signaling molecules related to integrins, and *Pecam1,* was relatively low in dormant HSCs (Fig. 6a), suggesting that impaired actin polymerization suppressed macrophage fragment adhesion on dormant HSCs. Third, live CD115^low^F4/80^+^ BM macrophages highly expressed phosphatidylserine (PS) compared to neutrophils and CD115^high^F4/80^+^ macrophages (Fig. 6b). An *in vitro* experiment with LT-HSCs revealed that masking PS by treatment with annexin V significantly inhibited the adhesion of apoptotic cell fragments (Fig. 6c), suggesting PS receptor-mediated macrophage fragment adhesion. On the other hand, dormant HSCs showed a relatively weak expression of PS receptors, such as *Itgav*, *Itgb3* and *Cd93* (Fig. 6a), possibly resulting in the impaired adhesion of macrophage fragments.

**Fig. 6:**
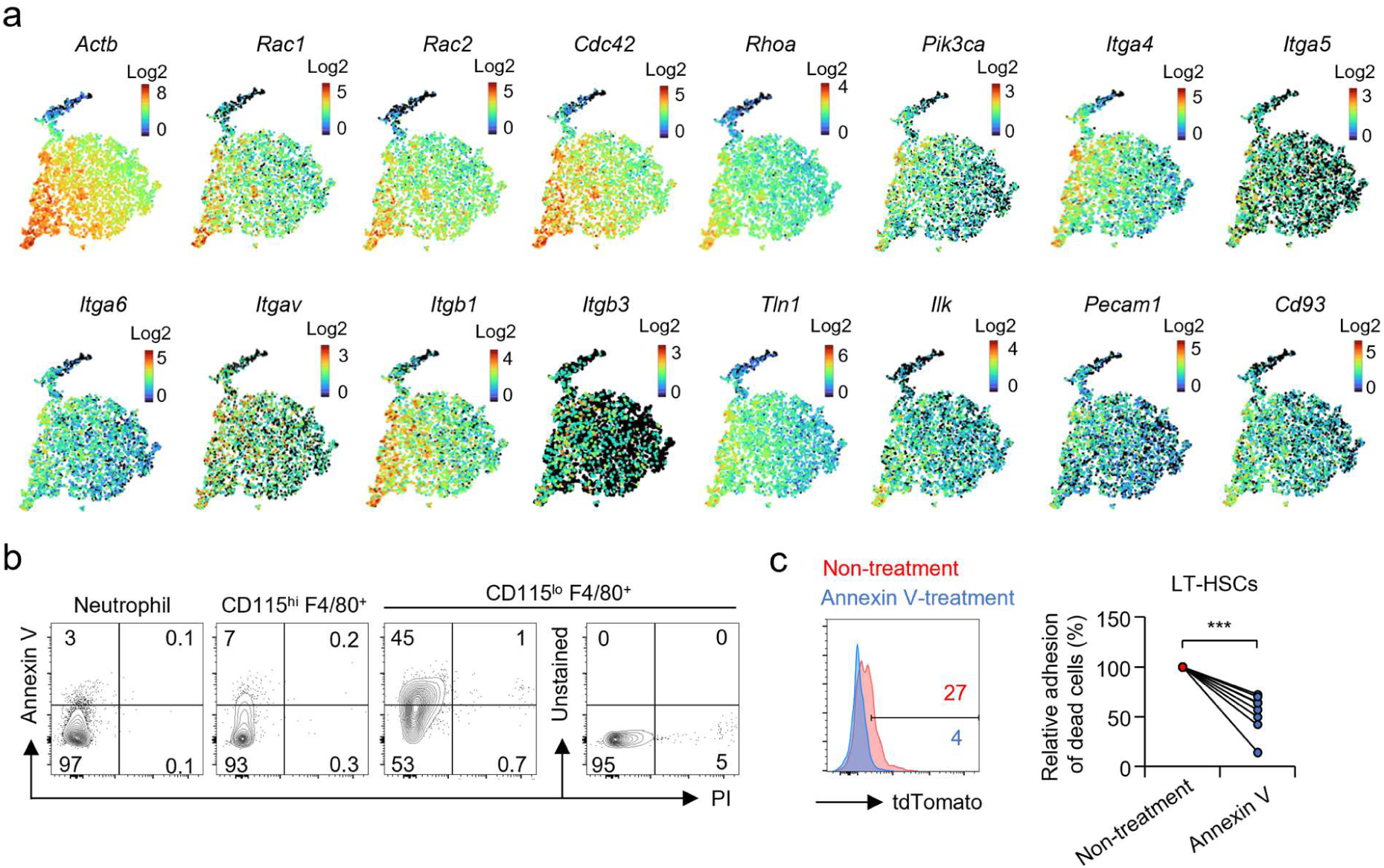
Exploring molecules that mediate the attachment of macrophage fragments to HSCs. **a** Expression of genes related to cell adhesion, cell-cell interactions and the recognition of dead cells on LT-HSCs. **b** Annexin V staining for neutrophils, and for CD115^+^F4/80^+^ and CD115^low^F4/80^+^ macrophages. **c** Inhibition of the attachment of dead cell fragments on the surface of LT-HSCs by treatment with annexin V. HSPCs obtained from *CAG-EGFP* mice were cultured with apoptotic tdTomato^high^ c-kit^-^ splenocytes in the presence or absence of annexin V (5 μg/ml) for 5 hours. The expression of tdTomato in EGFP^+^ LT-HSCs was examined by FACS. A representative FACS plot for LT-HSCs and statistical analysis are shown in the left and right panels, respectively. n=8 per group. ***p<0.001. Student’s t-test was used [c]. Data are representative of two independent experiments (b) or were pooled from three independent experiments (c).

In summary, the selective adhesion of macrophage fragments to proliferative HSCs, but not to dormant HSCs is mediated by multiple mechanisms, including CD169/CD43, actin polymerization, and PS receptors.

### F4/80^low^ HSCs retain considerable stemness during inflammation and aging

The lack of macrophage fragment adhesion was a valuable indicator for detecting dormant HSCs under the steady state. We tested whether this scenario can be applicable under stress conditions. Aging is considered a state of physiological stress and causes a variety of transformations in HSCs, namely reductions of self-renewal and regenerative capacities and myeloid biased differentiation^42^. Thus, we examined the validity of our method in aged mice. CITE-seq analysis was performed on LT-HSCs from 21-month-old WT mice (pink in Fig. 7a) and compared with those from 8-week-old mice (blue in Fig. 7a). A previous study reported that CD86^-^ HSCs with myeloid-biased potential emerged in aged mice^43^. Consistent with that report, CD86^-^ HSCs with an impaired reconstitution capacity increased considerably with age (Supplementary Fig. 7a, b), suggesting that identifying dormant HSCs is expected to be more difficult in the presence of CD86^-^ HSCs in aging mice. Thus, we performed CITE-seq analysis on CD86^+^ LT-HSCs and identified CD34^low^F4/80^low^CD169^low^ HSCs in aged mice (Fig. 7b c). Like young mice, the F4/80^low^ HSCs of aged mice showed a higher expression of Sca-1 and a lower expression of c-kit and CD48 compared to F4/80^high^ HSCs (Fig. 7d). In addition, the expression levels of *Actb*, β-actin rearrangement-associated genes, and cell-cycle-associated genes (shown in red in Fig. 5e) were low in the F4/80^low^ HSCs of aged mice (Fig. 7e and Supplementary Fig. 7c), suggesting the dormant state in aged F4/80^low^ HSCs. The number of F4/80^low^ HSCs was not significantly changed (Fig. 7f), and the expression levels of most genes related to dormancy were not reduced in aged F4/80^low^ HSCs, except for the downregulation of *Meg3* and *Igf2bp*, compared to young F4/80^low^ HSCs (Fig. 7g and Supplementary Fig. 7d). Furthermore, the expression of the growth-promoting proto-oncogene *Myc* was increased (Fig. 7g). In this context, the downregulation of *Meg3* and *Igf2bp2*, and the upregulation of *Myc*, occurred as early as 13 months of age in F4/80^low^ HSCs (Fig. 7h-k), despite the low expression of entire cell cycle-associated genes (Fig. 7l). We further tested whether our method is applicable to acute inflammation. To avoid contamination of myeloid progenitors in HSC gating by the upregulation of Sca-1 and to accurately identify LT-HSCs under inflammatory conditions, we used CD86 as an alternative marker for Sca-1, as previously reported^44^. CITE-seq analysis of LT-HSCs obtained from WT mice 24 hours after LPS-treatment revealed the existence of F4/80^low^CD169^low^ HSCs (Supplementary Fig. 7e, f), which showed lower expression of c-kit, CD48, CD34, and cell-cycle associated genes even during acute inflammation (Supplementary Fig. 7g, h). In addition, F4/80^low^ HSCs expressed *Meg3* and *Igf2bp2*, albeit somewhat attenuated compared to naïve mice (Supplementary Fig. 7i).

**Fig. 7:**
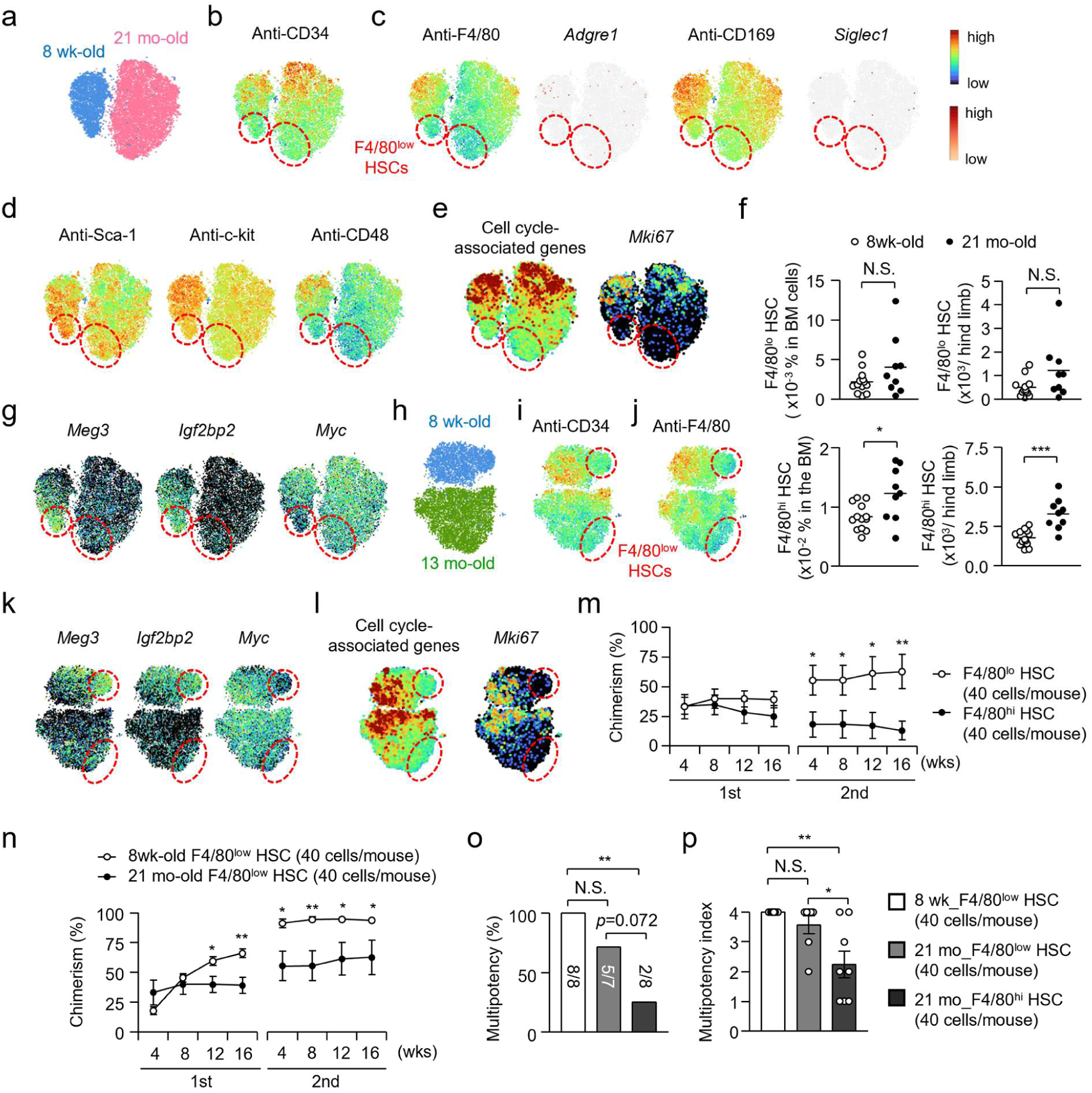
Identification of dormant HSCs by detecting macrophage fragments during biological stresses. **a-e, g** CITE-seq analysis of LT-HSCs obtained from 8 week-old (blue in a) and 21 month-old (pink in a) naïve WT mice (g-m). Expression levels of cell surface CD34 (b), cell surface F4/80 and/or CD169 and genes encoding those proteins (c), cell surface HSC markers (d), cell-cycle-associated genes (e), dormancy-related genes (g) and cell surface CD41 were examined in LT-HSCs. **f** Frequencies and numbers of F4/80^low^ HSCs in the BM of 8 week-old or 21 month-old mice. **h-l** CITE-seq analysis for LT-HSCs obtained from 8 week-old (blue in h) and 13 month-old (green in h) naïve WT mice. Expression levels of cell surface CD34 (i) and F4/80 (j), dormancy-related genes (k) and cell-cycle-associated genes (l). **m** Chimerism of donor cells derived from F4/80^low^ HSCs or F4/80^high^ HSCs obtained from aged mice. F4/80^low^ or F4/80^high^ HSCs (40 cells each) obtained from 21-month old mice were intravenously transplanted into lethally irradiated SJL mice with EGFP^+^ rescue BM cells (1 x 10^5^). n=7 for F4/80^low^ HSCs, n=8 for F4/80^high^ HSCs. **n** Chimerism of donor cells derived from F4/80^low^ HSCs. F4/80^low^ HSCs (40 cells) were obtained from 8 week-old or 21 month-old mice and were then transplanted into SJL mice with 10^5^ EGFP-expressing BM cells. n=8 for 8 week-old, n=7 for 21 month-old mice. **o, p** F4/80^low^ or F4/80^high^ HSCs obtained from 8 week-old or 21 month-old mice were transplanted into lethally irradiated recipients. The frequency of mice in which 4 lineages (B cells, T cells, neutrophils and monocytes) were reconstituted (o) and multipotency index, numbers of reconstituted lineages (p), were examined at 16 weeks after the 2^nd^ transplantation. The maximum score of the index is 4 (B cells, T cells, neutrophils and monocytes). n=8 for F4/80^low^ HSCs obtained from 8 week-old mice and F4/80^high^ HSCs obtained from 21 month-old mice, n=7 for F4/80^low^ HSCs obtained from 21 month-old mice. Error bars, SEM. *p<0.05, **p<0.01. Student’s t-test [f, m, n], χ^2^ statistics [o] and one-way ANOVA [p] were used. Data were pooled from two (n-p) or three (f) independent experiments.

Using cell cycle-associated genes obtained from CITE-seq analysis (Fig. 5e) as an indicator of dormant HSCs, we also evaluated the suitability of F4/80 and other known cell surface markers for dormant HSCs under aging and LPS-induced inflammation. Among them, F4/80^low^ is one of the most suitable markers for identifying dormant HSCs under such stress conditions (Supplementary Fig. 7j, k). In both 13- and 21-month-old mice, known dormant HSC markers^5, 6, 7, 8, 9, 10^ differed from the expression patterns of cell cycle-related genes (Supplementary Fig. 7k). In contrast, F4/80^low^ HSCs corresponded to HSCs with low expression of cell cycle-associated genes (Supplementary Fig. 7k), implying that they are dormant HSCs. We also observed that in aged mice, F4/80^low^ dormant HSCs appeared to differ from recently reported ‘young-like HSCs’ that weakly express aged HSC marker genes, such as *Clu, Clca3a1, Ramp2, Sbspon, and Gpr183*^45, 46, 47, 48^ (Supplementary Fig. 7k, l). Collectively, the lack of macrophage fragment adhesion appears to be a valuable indicator for dormant HSCs in these stress conditions.

To verify whether the F4/80^low^ HSCs in aged mice meet the functional criteria of dormant HSCs, we separately transplanted F4/80^low^ HSCs and F4/80^high^ HSCs from aged mice into lethally irradiated recipient mice. Compared to F4/80^high^ HSCs, the chimerism of F4/80^low^ HSC-derived donor cells was significantly higher after the second transplantation (Fig. 7m), indicating that the hierarchy between F4/80^low^ and F4/80^high^ HSCs was maintained in aged mice. To directly compare the stemness of F4/80^low^ HSCs between young and aged mice, we isolated F4/80^low^ HSCs from young and from aged mice and separately transplanted them into lethally irradiated recipient mice. The chimerism of aged F4/80^low^ HSC-derived cells was lower than that of young F4/80^low^ HSC-derived cells in the later phase of the 1^st^ and the entire 2^nd^ transplantation (Fig. 7n), indicating that aging somewhat reduced the BM-reconstitution ability of F4/80^low^ HSCs, which may be due to the altered expression of a few genes associated with dormancy and growth inhibition (Fig. 7g and Supplementary Fig. 7d). In contrast, F4/80^low^ HSCs from aged mice maintained a comparable level of multipotency as F4/80^low^ HSCs from young mice (Fig. 7o, p and Supplementary Fig. 7m, n), suggesting that F4/80^low^ HSCs retain considerable stemness in aged mice.

Thus, our novel method, based on macrophage fragment attachment, enabled the easy identification of dormant HSCs during all life stages and their direct comparison between naïve and stress conditions.

## Discussion

The HSC fraction is heterogeneous and contains biased HSCs, proliferative HSCs, dormant HSCs and cells without stemness^1, 2, 49^. The heterogeneity of HSCs can complicate the interpretation of experimental results, especially when comparing different conditions, e.g., steady state vs. stress conditions or WT mice vs. genetically modified mice, because the results may reflect a change of cellular composition in HSC fractions rather than an intrinsic alteration of HSCs. In this study, we developed a novel and simple method to identify proliferative and dormant HSCs separately by detecting macrophage fragment adhesion. Using this method, we demonstrated that HSCs without macrophage fragment attachment correspond to dormant HSCs, exhibiting delayed BM reconstitution, decreased long-term proliferative activity, the restricted expression of proliferation-related genes, and the enrichment of dormancy-related genes. Conveniently, this method is readily applicable to a variety of studies because F4/80 can be used as a marker and does not require additional treatment of HSCs. In this context, the dormancy of F4/80^low^ HSCs, characterized by quiescent cell cycle and prominent stemness, was largely maintained under conditions of inflammation and aging, suggesting that this method may enable a more accurate analysis of HSCs under biological stresses. In contrast, known markers for dormant HSCs did not correspond to HSCs with low expression of cell-cycle associated genes under such stress conditions. Notably, F4/80^low^ dormant HSCs contained a significant number of cells expressing aged HSC marker genes (Supplementary Fig. 7k, l), which is the opposite characteristic of ‘young-like HSCs’ ^45, 46, 47, 48^. In addition, ‘young-like HSCs’ exhibit a similar cell-cycle status to aged HSCs^45, 47^. Thus, the lack of macrophage fragment adhesion appeared to be a benchmark of dormant HSCs but not ‘young-like HSCs’ in aged mice.

A recent report suggested that HSCs acquire CXCR4 from macrophages through trogocytosis, and HSCs that have undergone trogocytosis express a macrophage marker F4/80^16^. That study also suggested that F4/80^-^ HSCs have a higher BM reconstitution capacity than F4/80^+^ HSCs and selectively mobilize to the periphery upon the injection of G-CSF, a CXCR4 inhibitor, and following macrophage depletion^16^, suggesting that HSCs with poor BM reconstitution capacity preferentially remain in the BM. This scenario seems unlikely because dormant HSCs play a physiological role as a reserve pool of HSCs and thus possess limited mobility^24, 50, 51^. In addition, if macrophage-derived CXCR4 on F4/80^+^ HSPCs is essential for their retention in the BM, treatment with a CXCR4 inhibitor should promote the emigration of F4/80^+^ HSPCs from the BM, leading to their emergence in the periphery. Instead, we proposed that HSCs have no chance to encounter and interact with macrophages and their fragments during the process of collecting mobilized HSCs from peripheral blood, because macrophages are absent in the blood. Consistently, while the HSPCs obtained from the blood of CXCR4 inhibitor-treated mice were F4/80^-^, most HSPCs became F4/80^+^ when mixed with a BM cell suspension that contains macrophages (Fig. 2e, f). Our IEM analysis further supported the attachment of macrophage fragments rather than trogocytosis (Fig. 2k and Supplementary Fig. 2k-n). Consistent with our findings, another report also suggested the presence of macrophage-derived molecules on HSCs, which reflected the attachment of macrophage fragments to the surface of HSCs during the cell preparation process *in vitro*^17^. In addition, macrophage-related molecules on HSCs could be detected at both the protein and mRNA levels^17^. However, the physiological relevance of macrophage-derived molecules on HSCs remained to be validated. We were unable to detect macrophage-related molecules, such as F4/80 and CD169, at the mRNA level (Fig. 5c and Supplementary Fig. 2f, 2i, 5), suggesting that those molecules primarily attach to HSCs and are dysfunctional. On the other hand, we detected TLR4 and CXCR4 at both the protein and mRNA levels (Fig. 3a, g), indicating that they are intrinsically expressed and functional in HSCs (Fig. 3a, b, g-h).

The adhesion of macrophage fragments to HSCs is mediated by multiple molecules related to actin-mediated activities, cell adhesion, cell-cell interactions, and dead cell recognition (Fig. 6 and Supplementary Fig. 6). We found that dormant HSCs weakly express the molecules involved in these processes, likely resulting in the impaired attachment of macrophage fragments, e.g. F4/80^lo^ HSCs. Our results also showed that CD115^low^ BM macrophages were predominant in the cell fragment compartment, but not in the dead cell fraction, implying that the BM macrophages are not prone to death but rather tend to fragment easily. EM analysis revealed that some large macrophage fragments retained the integrity of their membrane structures (Supplementary Fig. 6a, b) and showed the formation of clathrin-coated vesicles and unknown vesicles inside, but it is unclear whether the formation of those structures is related to the fragmentation of BM macrophages. The possibility that macrophage fragments may adhere to a small number of HSCs/HSPCs *in vivo* remains debatable. Our EM and confocal microscopy analysis implied that a few macrophage fragments might be phagocytosed by HSPCs (data not shown).

Given that macrophage fragments serve as a valuable indicator to easily distinguish between proliferative and dormant HSCs both in steady-state and in stress conditions without further manipulation of HSCs *in vitro*, it is expected that this unique method will make a significant contribution to our understanding of the bona fide hematopoietic response. Indeed, we directly compared dormant HSCs between young and aged mice, identifying *Igf2bp2*, *Meg3* and *Myc* as genes showing expression changes that reduce dormancy in the early stages of aging from a set of dormant HSC-specific genes. We also identified a gene set that is selectively expressed in dormant HSCs of naïve young mice as promising candidates regulating HSC stemness (Fig. 5e). Investigating the roles of these genes may lead to the discovery of novel mechanisms that maintain HSCs and their functions in the future.

## Methods

### Mice

All animal procedures were performed in accordance with the TMDU Animal Experiment Guidelines and were approved by the TMDU Institutional Animal Care and Use Committee. WT C57BL/6J (B6) and *CAG-EGFP* mice were obtained from Japan SLC (Hamamatsu, Japan); B6.SJL-ptprca (B6.SJL, Strain #: 4007) mice congenic at the CD45 locus (CD45.1^+^CD45.2^−^) were from Taconic (Germantown, NY, USA); *Rosa26-lsl-tdTomato* mice (Strain#: 007909), *Mrp8-Cre-ires-Gfp* mice (Strain#: 021614), *LysM-Cre* mice (Strain#: 004781), *Vav1-Cre* mice (Strain#: 035670) and *Tlr4^flox/flox^* mice (Strain#: 024872) were from Jackson Laboratories (Bar Harbor, ME, USA). *Cd169*-DTR mice were obtained from Dr. Masato Tanaka of Tokyo University of Pharmacy and Life Sciences. *Il7r*-Cre mice were previously generated by us^52^.

### Cell culture

HSPCs were cultured in RPMI1640 (FUJIFILM, Osaka, Japan, 18902025) containing 10% fetal bovine serum (FBS). All media were supplemented with penicillin-streptomycin (Nacalai tesque, Kyoto Japan, 09367-34), non-essential amino acids (Thermo Fisher Scientific, Waltham, MA, USA, 11140-050), sodium pyruvate (Thermo Fisher Scientific, 11360-070) and 2-Mercaptoethanol (Thermo Fisher Scientific, 21985-023).

### Cell identification by flow cytometry

Cells in adult mice were identified as follows; HSCs (CD34^-^ LT-HSC): CD34^-^Lin^-^c-kit^+^Sca-1^+^CD150^+^CD48^-^, HSPCs: Lin^-^c-kit^+^Sca-1^+^, LT-HSCs: Lin^-^c-kit^+^Sca-1^+^CD150^+^CD48^-^, ST-HSCs: Lin^-^c-kit^+^Sca-1^+^CD150^-^CD48^-^, MPP2s: Lin^-^c-kit^+^Sca-1^+^CD150^+^CD48^+^, MPP3s: Lin^-^c-kit^+^Sca-1^+^CD150^-^CD48^+^Flt3^-^, MPP4s: Lin^-^c-kit^+^Sca-1^+^CD150^-^CD48^+^Flt3^+^, CLPs: Lin^-^IL-7R^+^Flt3^+^CD86^+^, GMPs: Lin^-^c-kit^+^Sca-1^-^CD150^-^CD34^+^CD16/32^+^, CMPs: Lin^-^c-kit^+^Sca-1^-^CD150^-^CD34^+^CD16/32^-^, MEPs: Lin^-^c-kit^+^Sca-1^-^CD150^+^, LMPPs: Lin^-^c-kit^+^Sca-1^+^CD150^-^CD48^+^Flt3^+^VCAM-1^-^, GMLPs: Lin^-^c-kit^+^Sca-1^+^CD150^-^CD48^+^Flt3^+^VCAM-1^+^, neutrophils: CD3^-^CD19^-^c-kit^-^NK1.1^-^Gr-1^+^CD115^-^, monocytes: CD3^-^CD19^-^c-kit^-^NK1.1^-^Gr-1^+^CD115^+^, CD115^+^ macrophages: CD3^-^CD19^-^c-kit^-^NK1.1^-^Gr-1^-^CD115^+^F4/80^+^, CD115^-^ macrophages: CD3^-^CD19^-^c-kit^-^NK1.1^-^Gr-1^-^CD115^-^F4/80^+^, T cells: CD3^+^CD19^-^NK1.1^-^, B cells: CD3^-^CD19^+^, NKT cells: CD3^+^CD19^-^NK1.1^+^, NK cells: CD3^-^CD19^-^c-kit^-^NK1.1^+^, c-kit^+^ cells: CD3^-^CD19^-^c-kit^+^NK1.1^-^. Lineage markers for hematopoietic stem progenitors included CD3, CD19, B220, NK1.1, CD4, CD8, Gr-1, CD11b and TER119. For all cell lineages, propidium iodide (PI)^-^ live singlet cells were pre-gated before identification with specific markers.

### Cell sorting

For the sorting of HSCs or HSPCs, BM cells were stained with APC or PE/Cy7-conjugated anti-c-kit antibody and anti-APC or anti-Cy7 microbeads (Miltenyi Biotec, Bergisch Gladbach, Germany) and were pre-sorted by positive selection with AutoMACS pro (Miltenyi Biotec). For the sorting of F4/80^high^ or F4/80^low^ HSPCs, the pre-sorted cells were then stained with specific antibodies, and the Lin^-^c-kit^+^Sca-1^+^ fraction was isolated by a 1^st^ FACS sorting with FACS Aria III (BD Bioscience, Franklin Lakes, NJ, USA). F4/80^high^ or F4/80^low^ HSPCs were then sorted by a 2^nd^ FACS sorting to purify them thoroughly. For the sorting of F4/80^high^ or F4/80^low^ HSCs, the pre-sorted cells were stained with specific antibodies, and the CD34^-^Lin^-^c-kit^+^Sca-1^+^ fraction was isolated by a 1^st^ FACS sorting with FACS Aria III (BD Bioscience). F4/80^high^ or F4/80^low^ HSCs were then sorted by a 2^nd^ FACS sorting. When donor HSPCs were isolated from mixed BM chimera mice, the pre-sorted c-kit^+^ BM cells were stained with specific antibodies, and the CD45.1^-^CD45.2^+^EGFP^-^Lin^-^c-kit^+^Sca-1^+^ fraction was isolated by a 1^st^ FACS sorting. Then, EGFP^-^ donor HSPCs were isolated by a 2^nd^ FACS sorting.

### Generation of mixed BM chimera mice

To evaluate cell fragment attachment in LT-HSCs, tamoxifen (2 mg/mouse/day) was intraperitoneally injected into *Rosa26-lsl-tdTomato; Rosa26-CreERT2* mice for five consecutive days. Following that, BM cells harvested from *Rosa26-lsl-tdTomato; Rosa26-CreERT2* mice (CD45.2^+^) were mixed with BM cells obtained from *CAG-EGFP* mice (CD45.2^+^) at a 1:1 ratio and were transferred into lethally irradiated SJL mice (CD45.1^+^). Four weeks after the transplantation, the fluorescence of tdTomato in EGFP^high^ LT-HSCs or EGFP in tdTomato^high^ LT-HSCs was detected to determine the adhesion of cell fragments by FACS or confocal microscopy. To evaluate molecule acquisition from macrophages, BM cells obtained from SJL (CD45.1^+^) and *Tlr4^flox/flox^; Vav1-Cre* mice or CAG-EGFP mice (CD45.2^+^) were mixed at a 9:1 ratio and intravenously transplanted into lethally irradiated SJL mice (CD45.1^+^). Four weeks after the transplantation, CD45.1^-^CD45.2^+^ donor HSPCs or LT-HSCs were analyzed by flow cytometry. For some experiments, CD45.1^-^CD45.2^+^ donor HSPCs were sorted and stimulated with ultrapure LPS (tlrl-3pelps, 100 mg/ml, InvivoGen, San Diego, CA, USA) for 6 hours to examine the upregulation of inflammatory cytokine mRNAs such as *Il6*. To evaluate the loss of TLR4 from WT HSPCs, BM cells obtained from SJL (CD45.1^+^) and *Tlr4^flox/flox^; Vav1-Cre* mice (CD45.2^+^) were mixed at a 1:9 ratio and intravenously transplanted into lethally irradiated *Tlr4^flox/flox^; Vav1-Cre* mice (CD45.2^+^). Four weeks after the transplantation, CD45.1^-^CD45.2^+^ donor HSPCs or LT-HSCs were analyzed by flow cytometry. To evaluate *Tlr4* mRNA expression in donor HSCs, BM cells obtained from SJL (CD45.1^+^) and *Tlr4^flox/flox^; Vav1-Cre* mice (CD45.2^+^) were mixed at a 8:2 ratio and intravenously transplanted into lethally irradiated SJL mice (CD45.1^+^). Four weeks after the transplantation, CD45.1^-^CD45.2^+^ donor LSKs were isolated using flow cytometry, and the expression of *Tlr4* was examined by qPCR after 6 hours of stimulation with ultrapure LPS (100 μg/ml). To evaluate IL-6 production, total HSPCs were isolated from mixed BM chimera mice and were stimulated with ultrapure LPS (100 μg/ml) for 6 hours. The cells were treated with GolgiPlug (BD Biosciences, Franklin Lakes, NJ, USA) for the last 4 hours and fixed and permeabilized with BD Cytofix/Cytoperm™ Fixation/Permeabilization Kit (BD Biosciences), after which intracellular IL-6 was detected in WT or *Tlr4* cKO donor HSPCs and LT-HSCs by flow cytometry.

### Trypsin-EDTA treatment of HSPCs

HSPCs and LPMs were obtained from the BM and peritoneal cavity of naïve WT mice, respectively. The cells were incubated in the presence or absence of 0.03% trypsin-EDTA (Thermo Fisher Scientific) at 37 °C for 15 min, after which cell surface F4/80, Sca-1 or c-kit was detected in HSPCs and LT-HSCs by flow cytometry.

### Transwell migration assay

F4/80^low^ and F4/80^high^ HSPCs were sorted from the BM of WT mice by MACS sorting and subsequent FACS sorting. Transwells (3.0 μm pore size, 12 well plate, Costar, Boston, ME, USA) were pre-incubated with cell culture medium for 4 hours. Recombinant mouse CXCL12 (0, 20, 50, or 200 ng/ml, Biolegend, San Diego, CA, USA) was added in the lower chambers, and F4/80^low^ or F4/80^high^ HSPCs (6 x 10^4^ cells) were applied to the upper chambers. After 24 hours of culture, the numbers of cells that had migrated into the medium of the lower chambers were counted. Specific migration toward CXCL12 was calculated by subtracting “migrated cell number without CXCL12” from each migrated cell number.

### Blockade of CD169 during BM cell preparation

BM cells were obtained from the left and right tibia by flushing with MACS buffer and MACS buffer containing neutralizing antibody for CD169 (R&D Systems, Minneapolis, MN, USA, 5 μg/ml), respectively. Cells were stained with specific antibodies in the absence or presence of neutralizing antibody for CD169 (5 μg/ml) on ice for 30 min. After washing with anti-CD169 antibody-containing or non-containing MACS buffer, cells were washed and resuspended in anti-CD169 antibody-containing or non-containing MACS buffer, and the expression of F4/80 in HSPCs was examined by FACS.

### Induction of HSPC emigration to blood

WT B6 mice were treated with eight subcutaneous injections of G-CSF (125 μg/kg, Biolegend) every 12 hours. Three hours after the last administration, blood samples were collected. For treatment with the CXCR4 inhibitor, blood samples were obtained 1 hour after a single intraperitoneal injection of a CXCR4 inhibitor (5 mg/kg, AMD3100, Sigma-Aldrich, St Louis, MO, USA). Leukocytes obtained from the blood were mixed with a BM cell suspension from SJL mice and were stained with antibodies for analysis by FACS.

### Electron microscopy

Lin^-^ BM cells isolated from naïve WT mice were stained with biotinylated anti-F4/80 antibody and antibodies for cell surface markers. After washing, cells were incubated in the presence of Streptavidin-gold conjugate (15 nm, 0.015 mg/ml) for 30 min on ice. The HSPCs were then fixed by a conventional fixation method (1.6% paraformaldehyde and 3% glutaraldehyde in 0.1 mol/L phosphate buffer at pH 7.4, followed by an aqueous solution of 1% osmium tetroxide). Fixed samples were embedded in Epon 812, and thin sections (70–80 nm) were then cut and stained with uranyl acetate and lead citrate for observation using a Jeol-1010 electron microscope (Jeol, Tokyo, Japan) at 80 kV^53^.

### Proliferation assay

For DAPI staining, BM cells from naïve WT mice were stained with antibodies and Lin^-^ cells were briefly sorted by MACS using PE/Cy5-conjugated Lin antibodies and anti-Cy5 microbeads. The sorted Lin^-^ cells were fixed and permeabilized using a BD Cytofix/Cytoperm™ Fixation/Permeabilization Kit (BD Biosciences). After washing with the BD Cytoperm Wash buffer, cells were stained with DAPI (2 μg/ml in BD Cytoperm Wash buffer). After incubation on ice for 20 min, cells were centrifuged and re-suspended in MACS buffer for analysis with FACS. For BrdU staining, WT B6 mice were given BrdU in their drinking water (0.8 mg/ml) for 2 weeks. c-kit^+^ cells were briefly isolated from BM cells by MACS with a PE/Cy7-conjugated c-kit antibody and anti-Cy7 microbeads (Miltenyi Biotec) and they were stained with antibodies to cell surface markers. Cells were fixed and permeabilized using a BD Pharmingen™ APC BrDU Kit (BD Biosciences) and stained with an APC-conjugated anti-BrdU antibody. Cells were then centrifuged and re-suspended in MACS buffer for analysis with FACS.

### Confocal microscopy

LT-HSCs were sorted from the BM of CAG-EGFP; LysM-cre; Rosa26-lsl-tdTomato mice or mixed BM chimera mice generated with BM cells obtained from *CAG-EGFP* mice and *Vav1-Cre; Rosa26-lsl-tdTomato* mice and fixed with 4% paraformaldehyde. Cells were embedded with ProLong^TM^ Gold antifade reagent with DAPI (Thermo Fisher Scientific) and then analyzed using confocal microscopy (TCS SP8, Leica Microsystems Gmbh, Wetzlar, Germany).

### Real-time PCR

For evaluation of *Il6* expression in HSPCs, cells were stimulated with ultrapure LPS (100 μg/ml)(tlrl-3pelps, Invivogen) for 6 hours. Total mRNAs were isolated using a RNeasy micro kit (Qiagen, Hilden, Germany) and were then reverse-transcribed to cDNAs with FastGeneTM Scriptase II (Nippon Genetics, Tokyo, Japan) after which gene expression levels were determined using a Light Cycler 480 and SYBR Green I Master (04707516001, Roche Diagnostics, Basel, Switzerland). The values were normalized according to the expression of β-actin. Specific primers used for real-time PCR are as follows: *Tlr4* forward: ATGCATGGATCAGAAACTCAGCAA, *Tlr4*reverse: AAACTTCCTGGGGAAAAACTCTGG, *Il6*forward: GAGGATACCACTCCCAACAGACC, *Il6* reverse: AAGTGCATCATCGTTGTTCATACA, *Adgre1* forward: TGACTCACCTTGTGGTCCTAA, *Adgre1* reverse: CTTCCCAGAATCCAGTCTTTCC, *Cxcr4* forward: TCAGTGGCTGACCTCCTCTT, *Cxcr4* reverse: CTTGGCCTTTGACTGTTGGT, *Actb* forward: TGTTACCAACTGGGACGACA, *Actb* reverse: CTGGGTCATCTTTTCACGGT.

### *In vitro* evaluation of the attachment of dead cell fragments to HSCs

tdTomato^high^c-kit^-^ apoptotic cells were prepared by culturing splenocytes obtained from *Vav1-cre; Rosa26-lsl-tdTomato* mice with staurosporine (1 μM, Sigma-Aldrich) overnight. After washing, HSPCs isolated from *CAG-EGFP* mice were co-cultured with the apoptotic cells at a ratio of 1:10 for 5-6 hours in fresh IMDM media with 10% FBS in the presence or absence of purified recombinant annexin V (5 µg/ml, BD Biosciences). The attachment of apoptotic cells was evaluated by FACS analysis.

### RNA-sequencing

RNA sequence library preparation, sequencing, mapping, gene expression, and gene ontology (GO) enrichment analysis were performed using DNAFORM (Yokohama, Kanagawa, Japan). The quality of total RNAs was assessed using a Bioanalyzer (Agilent) to ensure that the RIN (RNA integrity number) was over 7.0. Double-stranded cDNA libraries (RNA-seq libraries) were prepared using a SMART-Seq Stranded Kit (Clontech, Mountain View, CA, USA, 634442) and a DNBSEQ MGIEasy Universal Library Conversion Kit (MGI Tech, Shenzhen, Guangdong, China) according to the manufacturer’s instructions. RNA-seq libraries were sequenced using paired-end reads (150 nt of read1 and read2) on a DNBSEQ-G400RS instrument (MGI Tech). The raw reads obtained were trimmed and quality-filtered using Trim Galore! (version 0.6.4_dev), Trimmomatic (version 0.39) and cutadapt (version 2.10) software. Trimmed reads were mapped to the mouse GRCm39 (mm39) genome using STAR (version 2.7.9a). Reads on annotated genes were counted using featureCounts (version 2.0.1). FPKM values were calculated from mapped reads by normalizing to total counts and transcripts. Differentially expressed genes were detected using the DESeq2 package (version 1.20.0). The list of differentially expressed genes detected by DESeq2 (basemean > 5 and fold-change < 0.25, or basemean > 5 and fold-change > 4) was used for GO enrichment analysis using the clusterProfiler package^54^.

### CITE-seq analysis

Lin (CD3, CD4, CD8, CD19, B220, NK1.1, TER119, Gr-1)^-^ cells were isolated from the BM cells of naïve 8 week-old WT mice, naïve 13 or 21 month-old WT mice or LPS (5 mg/kg)-treated 8 week-old WT mice by negative MACS sorting with Cy5-conjugated anti-Lin antibody and anti-Cy5 microbeads. LT-HSCs (Lin^-^Sca-1^+^c-kit^+^CD86^+^CD48^-^CD150^+^ BM cells) were then isolated by performing FACS sorting twice and stained with TotalSeq™-B Mouse Universal Cocktail, V1.0 (Biolegend). Cell counts and viability were measured using a hemacytometer with trypan blue staining. Approximately 10,000 cells with a 90% survival rate were applied to GEM (Gel Bead-in-Emulsion) generation using a Chromium Controller (10x Genomics), where individual cells were captured into single oil emulsions with reverse transcription reagent and a Gel Bead which contains barcoded oligonucleotides. The reverse transcription reaction enables individual cDNAs to be labeled by unique cell barcodes with unique molecular tags in each single cell emulsion. cDNA amplification followed by library construction for single-cell gene expression was performed using Chromium Single Cell 3’ Reagent Kits User Guide (v3.1 Chemistry) with Feature Barcoding technology for Cell Surface Protein (10x Genomics). A part of the cDNA was used for amplification and library preparation of antibody-derived fragments (A.K.A Cite-Seq). The resulting libraries were quantified by Qubit dsDNA Assay (ThermoFisher Scientific) and TapeStation D1000 ScreenTape (Agilent Technologies). The libraries were then pooled together at a ratio of 9:1 for gene expression and CITE-Seq and sequenced by a NovaSeq 6000 platform (Illumina) with 150 bp paired-end configuration, generating approximately 300M paired-end reads per sample. The sequence reads were further trimmed into Read1: 28 bp and Read2: 90 bp and processed into a Cell Ranger pipeline (10x Genomics), in which mapping, gene expression count, cell calling, and clustering analyses were performed. The single-cell gene expression and CITE-Seq analyses described above were conducted by GENEWIZ from Azenta Life Sciences (Tokyo, Japan).

### Statistical analysis

Statistical analyses were performed using Microsoft Excel or Prism software version 3 (GraphPad, San Diego, CA, USA). A two-tailed Student’s *t*-test was used for statistical analyses of two-group comparisons. Multigroup comparisons were performed using one-way analysis of variance (ANOVA) followed by the Tukey–Kramer multiple comparisons test. χ^2^ statistics were derived using the CHIDIST function of Microsoft Excel. P-values for correlations were calculated using the TDIST function of Microsoft Excel. The criterion of significance was set at *p* < 0.05. Results of biological replicates are expressed as means ± standard deviation (SD), and data of technical replicates are expressed as means ± standard error of the mean (SEM). Blinding or randomization of the groups was not performed, and no data were excluded. No statistical methods were used to estimate sample size.

## Data availability

All data are available in the main text or the supplementary materials. The RNA-seq and scRNA-seq data were deposited in the National Center for Biotechnology Information Gene Expression Omnibus with accession numbers GSExxxxxx, GSExxxxxx and GSExxxxxx.

## Acknowledgments

This work was supported by the Japan Society for the Promotion of Science (KAKENHI, Grant numbers 23K24151ZA and 18K19439, M.K.), the Japan Science and Technology Agency (PRESTO, Grant Number 11KA500235, M.K.), The Chemo-Sero-Therapeutic Research Institute Grant (KAKETSUKEN, M.K.), Multilayered Stress Diseases, Science Tokyo (T.O.), and Medical Research Center Initiative for High Depth Omics, Science Tokyo (T.O.). We thank H. Kamioka, Y. Watanabe and T. Akashi for their general assistance and M. Tanaka for providing *Cd169-DTR* mice.

## Author contributions

M. K. planned and performed the majority of experiments; Y. I. performed experiments to prove the attachment of macrophage fragments to HSCs; Y. Y. performed experiments for mixed chimera mice with *Tlr4*-deficient mice; S. A. performed experiments and provided suggestions for EM analysis; A. I. provided suggestions for experimental strategies for the analysis of macrophage fragment adhesion; M. K. and T. O. conceived of the project and wrote the manuscript.

