## Supplementary Figures 1-7 for "The lack of macrophage fragment adhesion is a benchmark of dormant hematopoietic stem cells throughout the lifespan"

### Supplementary Fig. 1

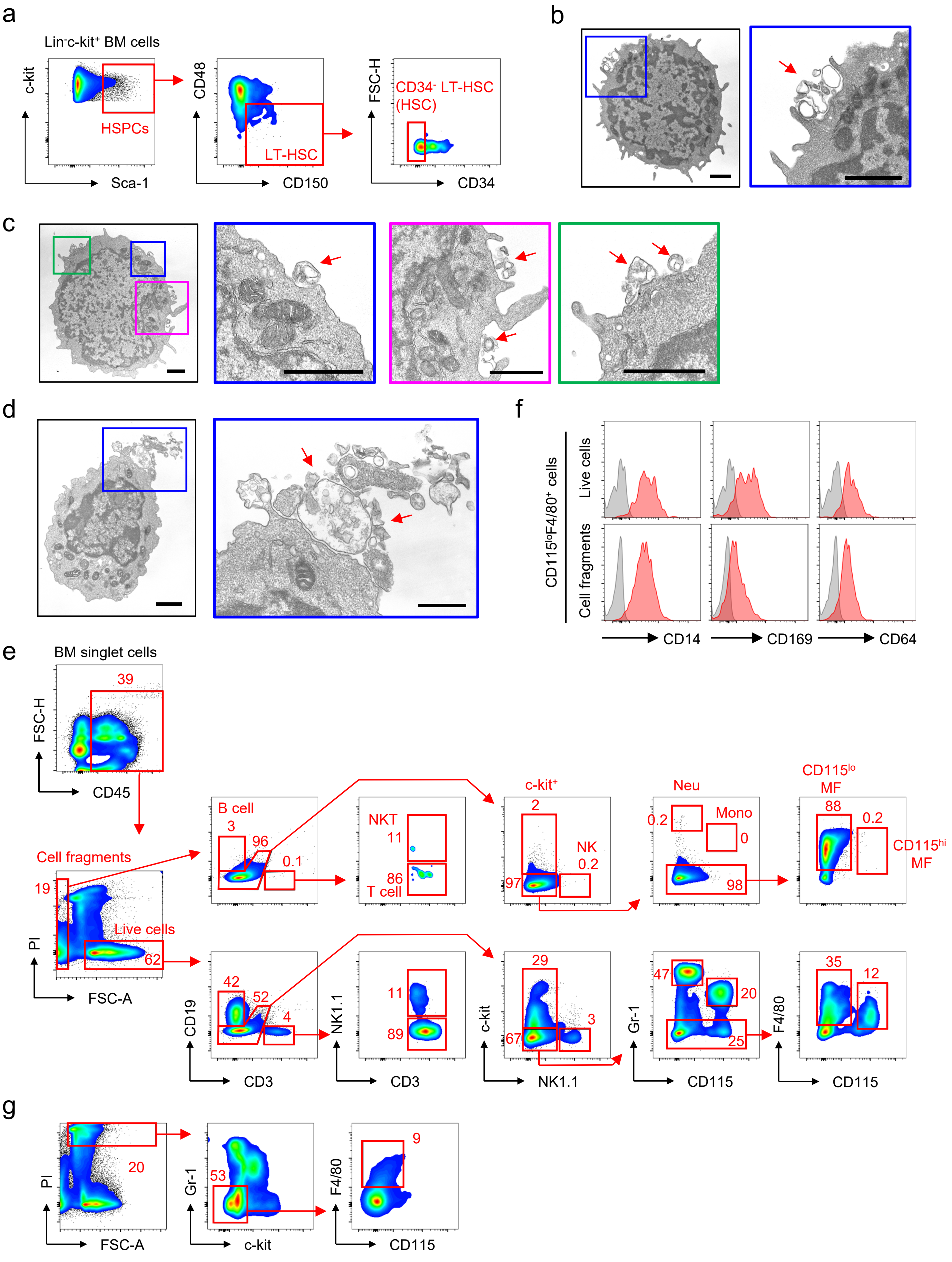

**Supplementary Fig. 1: Macrophage fragments in BM cell suspensions.**
**a** Gating strategy of HSPCs, LT-HSCs and HSCs. **b-d** EM analysis of HSPCs obtained from the BM of naïve WT mice. Red arrow indicates cell fragments attached to the surface of HSPCs. Scale bars indicate 1 μm. **e** Gating strategy for live cells and cell fragments in the BM. **f** Expression of macrophage markers in $CD115^{lo}F4/80^{+}$ cells in live cells and cell fragments in the BM. **g** Frequency of $CD115^{lo}F4/80^{+}$ cells in dead cells. Data are representative of two independent experiments (a-g).

### Supplementary Fig. 2

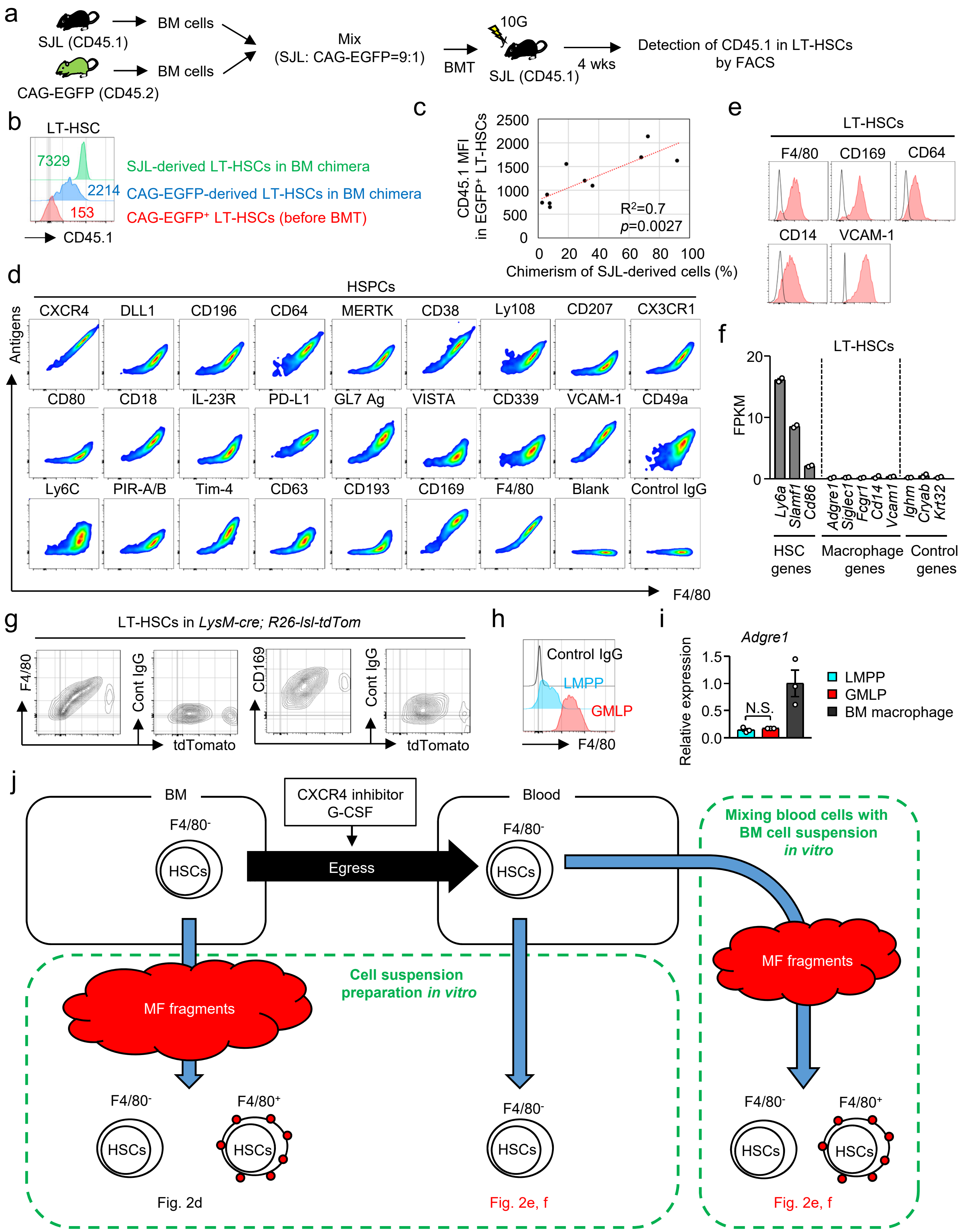

Supplementary Fig. 2 (continued)

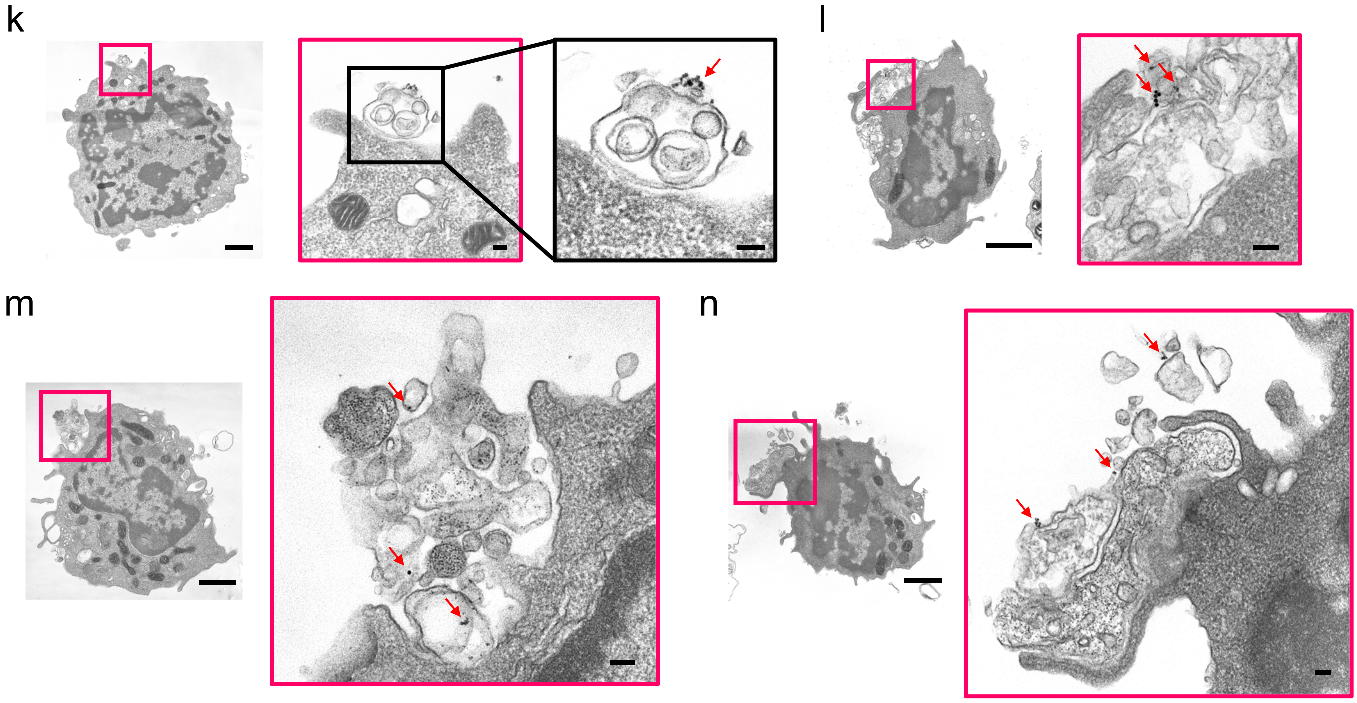


**Supplementary Fig. 2: Detection of macrophage-derived molecules on the surface of LT-HSCs.**

**a-c** BM cells obtained from SJL mice (CD45.1$^{+}$) were mixed with CAG-EGFP mice (CD45.2$^{+}$) at a 9:1 ratio (a, b) or at the indicated ratio (c). The mixed cells were then intravenously transferred into lethally irradiated SJL recipient mice. Before and 4 weeks after the transplantation, the expression levels of CD45.1 on the surface of EGFP$^{+}$ donor LT-HSCs were examined by flow cytometry. The numbers in the FACS plot indicate the mean fluorescence intensity (MFI) of CD45.1 (b). P-value was calculated using the TDIST function of Microsoft Excel (c). **d, e** The expression of F4/80 and the indicated molecules on the cell surface of HSPCs obtained from WT mice. **f** RNA-sequencing analysis of LT-HSCs obtained from naïve WT mice. Gene expression levels are shown by FPKM values. n=2. **g** Expression of tdTomato and F4/80 or CD169 in LT-HSCs obtained from *LysM-Cre; Rosa26-lsl-tdTomato* mice. **h** Expression of F4/80 on the surface of LMPPs and GMLPs in naïve WT mice. **i** Gene expression levels of *Adgre1* in LMPPs, GMLPs and BM macrophages in naïve WT mice evaluated by qPCR. **j** Schematic illustration of the mechanism by which HSCs acquire macrophage-derived cell surface molecules. **k-n** IEM of F4/80 on the surface of HSPCs. Red arrows indicate gold particles detecting F4/80. Scale bar in the leftmost panel in each figure indicates 1 µm and scale bars in the right and/or front panels indicate 0.1 µm. N.S.; not significant. One-way ANOVA was used [i]. Data are representative of two independent experiments (b, d, e, g, h) or were pooled from two (f) or three (c, i) independent experiments.

### Supplementary Fig. 3

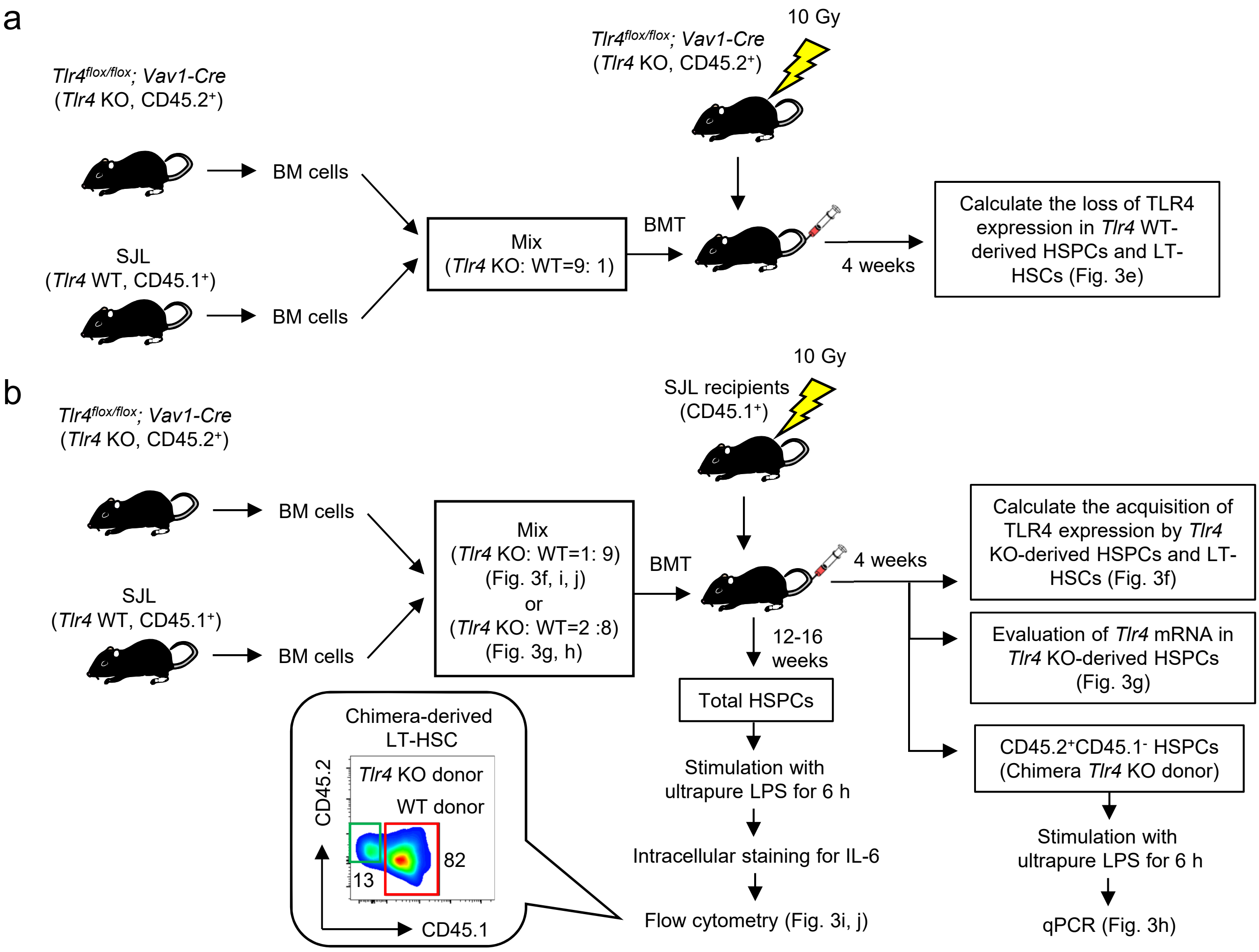


**Supplementary Fig. 3: Generation of mixed BM chimera mice.**
**a** Experimental strategy for Fig 3e. BM cells of *Tlr4* WT (SJL, CD45.1$^+$) mice and *Tlr4* KO (*Tlr4* $^{flox/flox}$; *Vav1*-cre, CD45.2$^+$) mice were mixed at a 1:9 ratio and then transferred into recipient *Tlr4* KO mice. The loss of TLR4 in SJL donor mice was calculated by comparing the TLR4 MFI in SJL donor mice between before and after transplantation. BMT: bone marrow transfer. **b** Experimental strategy for Fig. 3f-j. BM cells of *Tlr4* WT (CD45.1$^+$) mice and *Tlr4* KO mice (CD45.2$^+$) were mixed at 9:1 or 8:2 ratios and transferred into recipient SJL mice. The acquisition of TLR4 in *Tlr4* KO donor mice was calculated by comparing the TLR4 MFI between *Tlr4* KO donor mice and *Tlr4* WT donor mice. BMT: bone marrow transfer.

### Supplementary Fig. 4

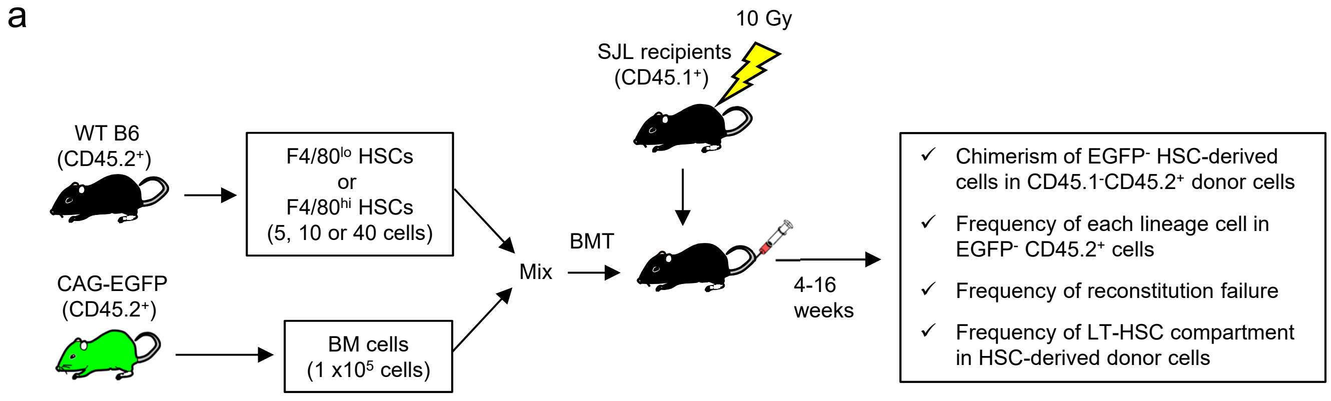


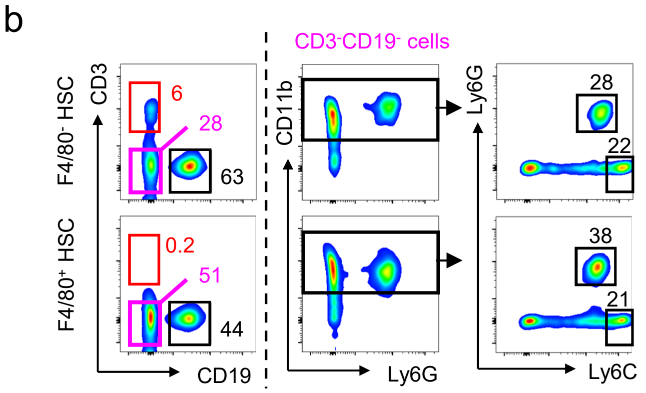


**Supplementary Fig. 4: Stemness of F4/80$^{lo}$ and F4/80$^{hi}$ HSCs.**
**a** Experimental strategy for Fig. 4c-k. The indicated numbers of F4/80$^{lo}$ or F4/80$^{hi}$ HSCs were mixed with EGFP$^{+}$ BM cells (1 x 10$^{5}$) and intravenously injected into lethally irradiated SJL mice. Four to sixteen weeks later, the chimerism of EGFP$^{-}$ cells in donor cells and/or frequencies of each lineage cell were examined in the mixed BM chimera mice. **b** Gating strategy for T cells, B cells, monocytes and neutrophils in the blood. Data are representative of three independent experiments.

Supplementary Fig. 5

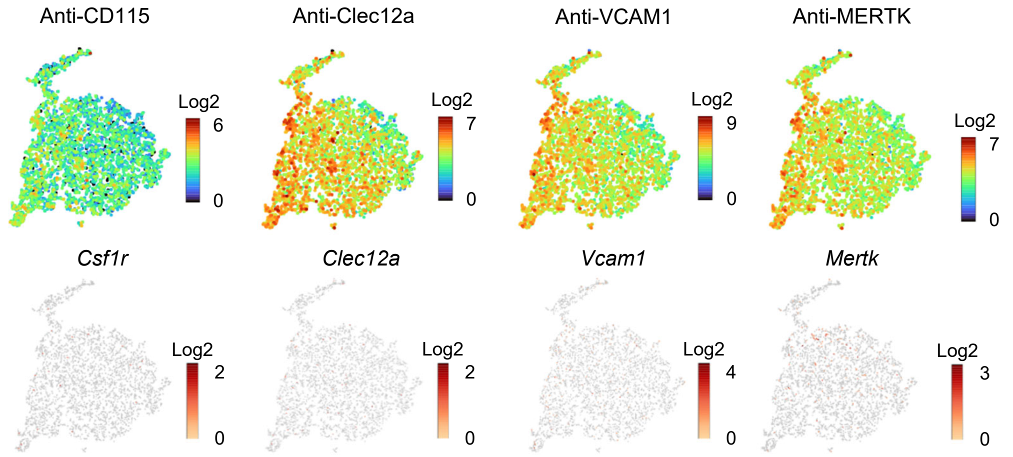


**Supplementary Fig. 5: Expression of macrophage-related molecules on F4/80$^{lo}$ HSCs.**
Cell surface expression of CD115, Clec12a, CD106 and MERTK and mRNA expression levels of *Csf1r*, *Clec12a*, *Vcam1* and *Mertk* in LT-HSCs.

Supplementary Fig. 6

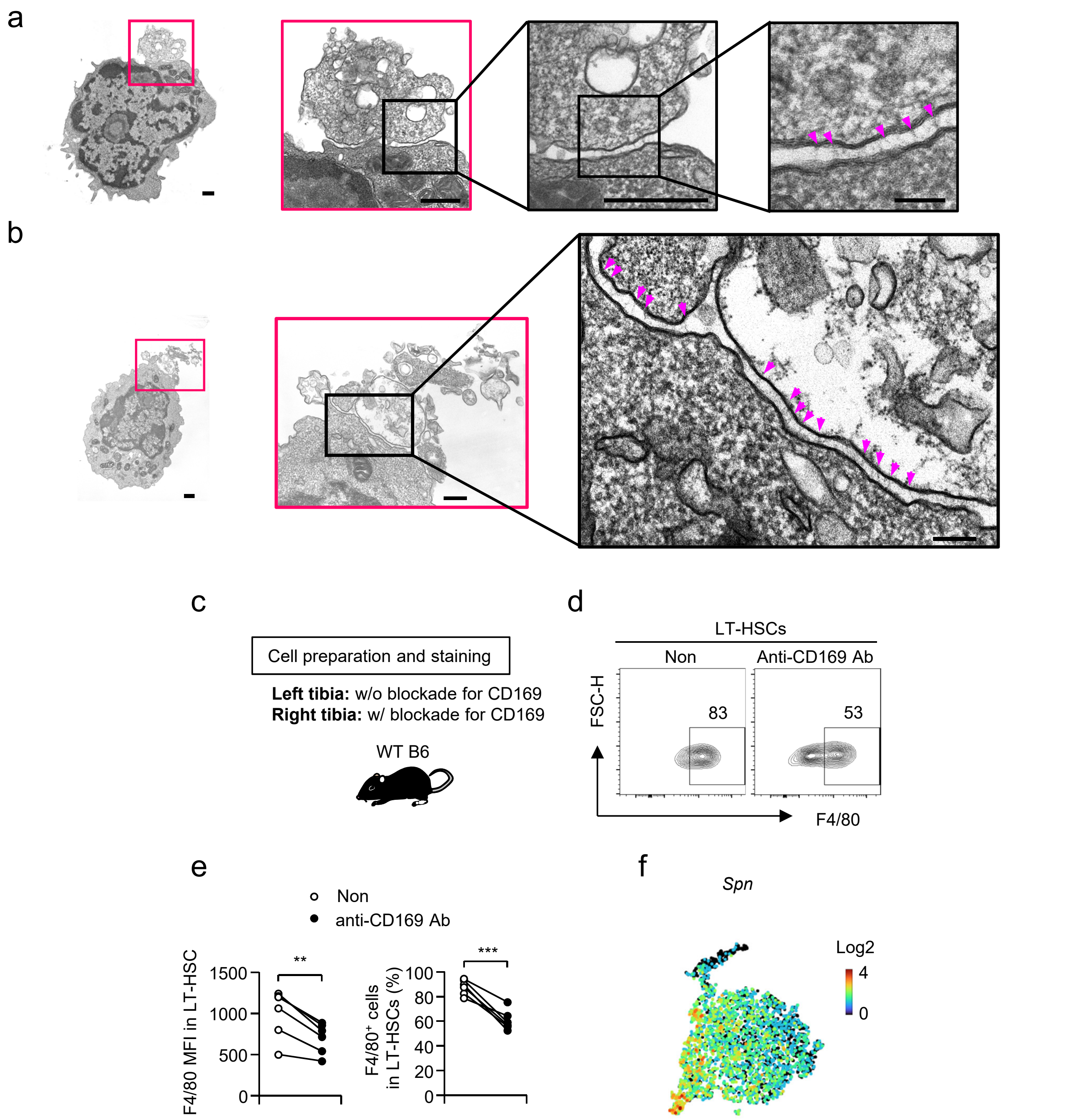


**Supplementary Fig. 6: Expression of *Spn*, a ligand for CD169, in F4/80$^{lo}$ HSCs.**
**a, b** EM of HSPCs obtained from the BM of naïve WT mice. Pink arrowheads indicate receptor-ligand interaction-like structures. The scale bar in the rightmost panel in (a, b) indicates 0.1 µm and scale bars in the other panels indicate 0.5 µm. The EM photos were shared with Fig. 1l and Supplementary Fig. 1d. **c** Experimental strategy for the preparation of BM cell suspension with blockade of CD169. **d, e** The expression of F4/80 in LT-HSCs prepared in the presence or absence of anti-CD169 antibody. Representative FACS plots are shown in (d) and the MFI of F4/80 and the frequency of F4/80$^{+}$ cells in LT-HSCs are shown in (e). **f** Expression of *Spn* in LT-HSCs. **p<0.01, ***p<0.001. Student's t-test was used. Data are representative of three pooled independent experiments (e).

### Supplementary Fig. 7

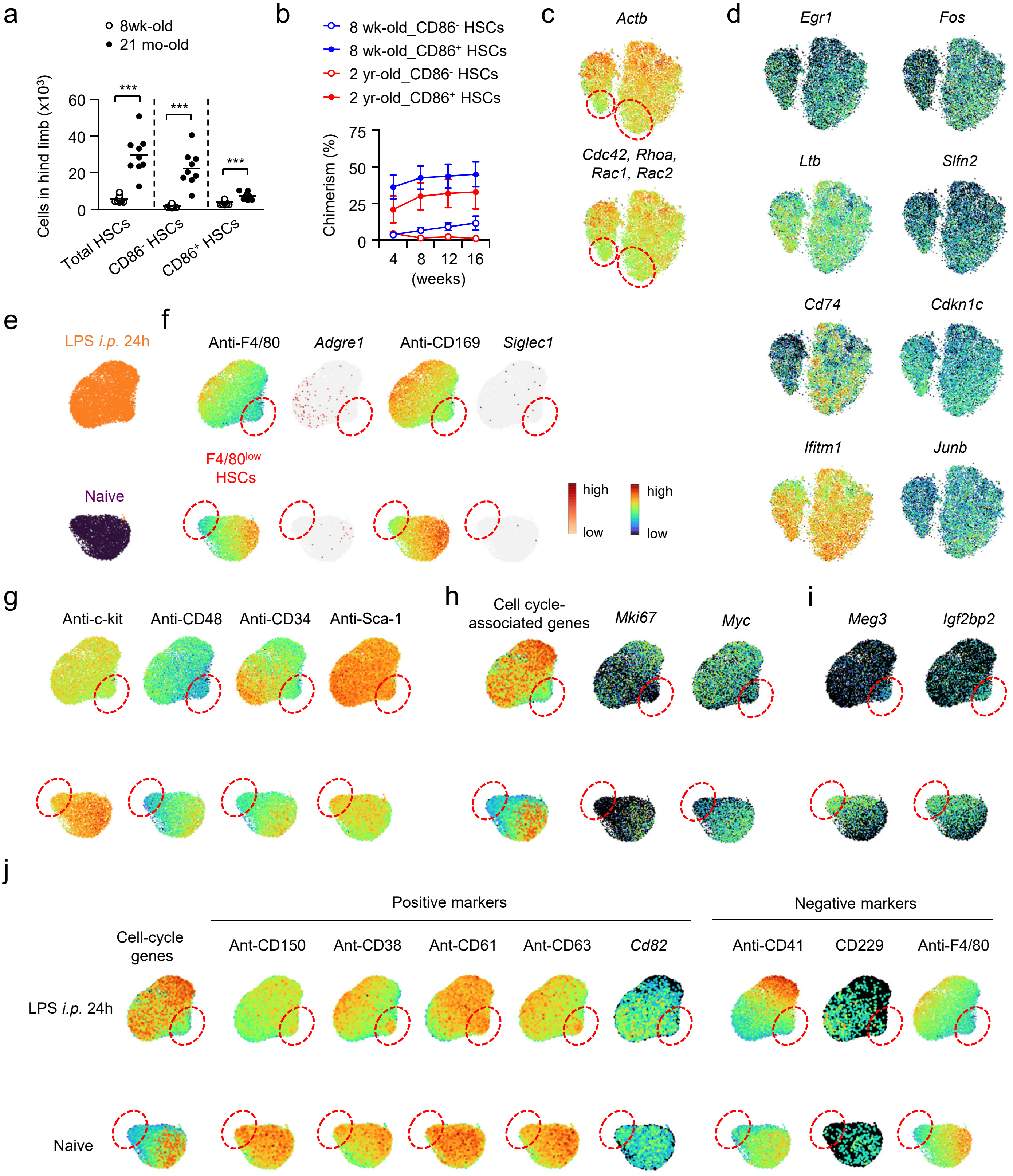

### Supplementary Fig. 7 (continued)

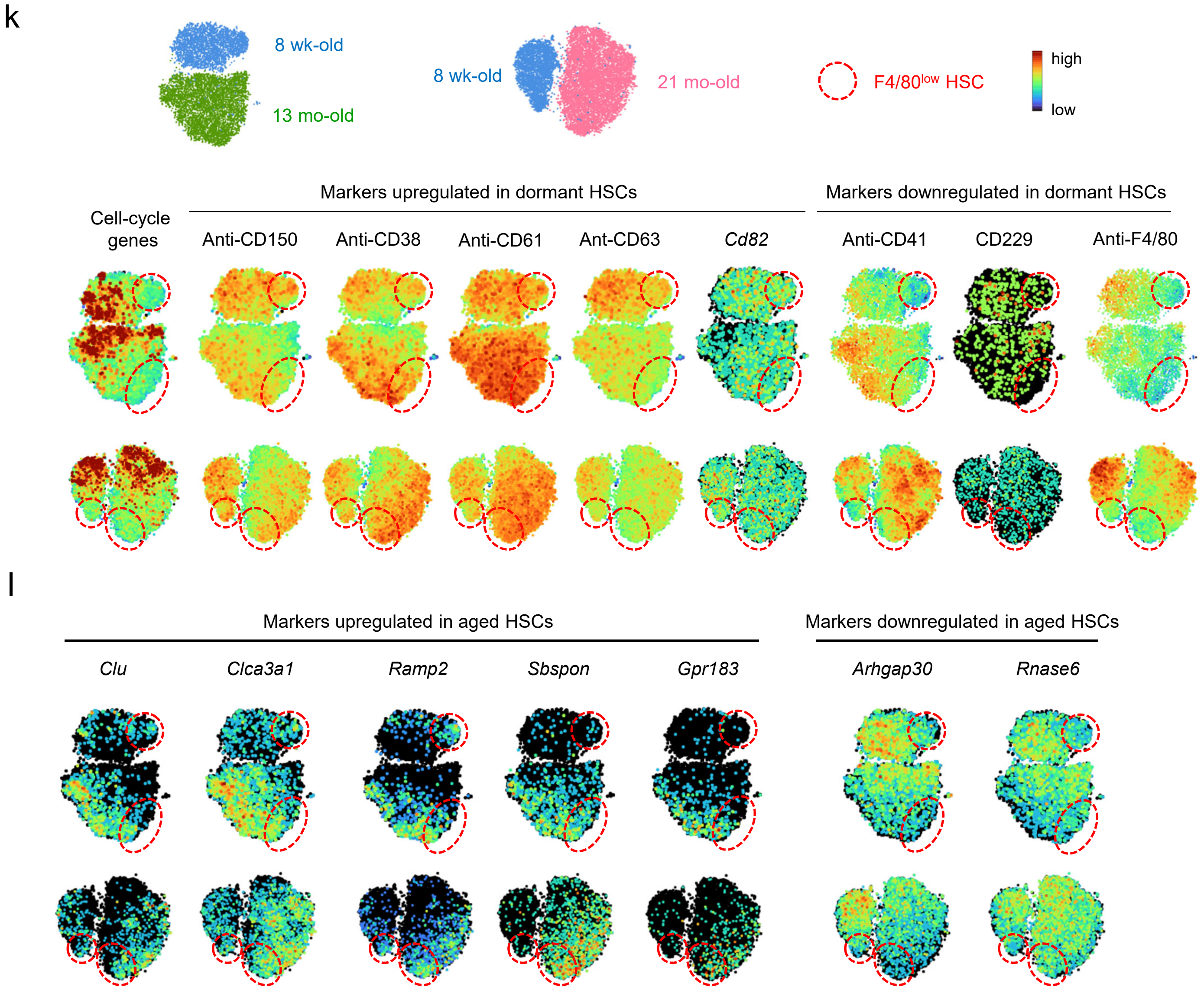

#### Supplementary Fig. 7 (continued)

m

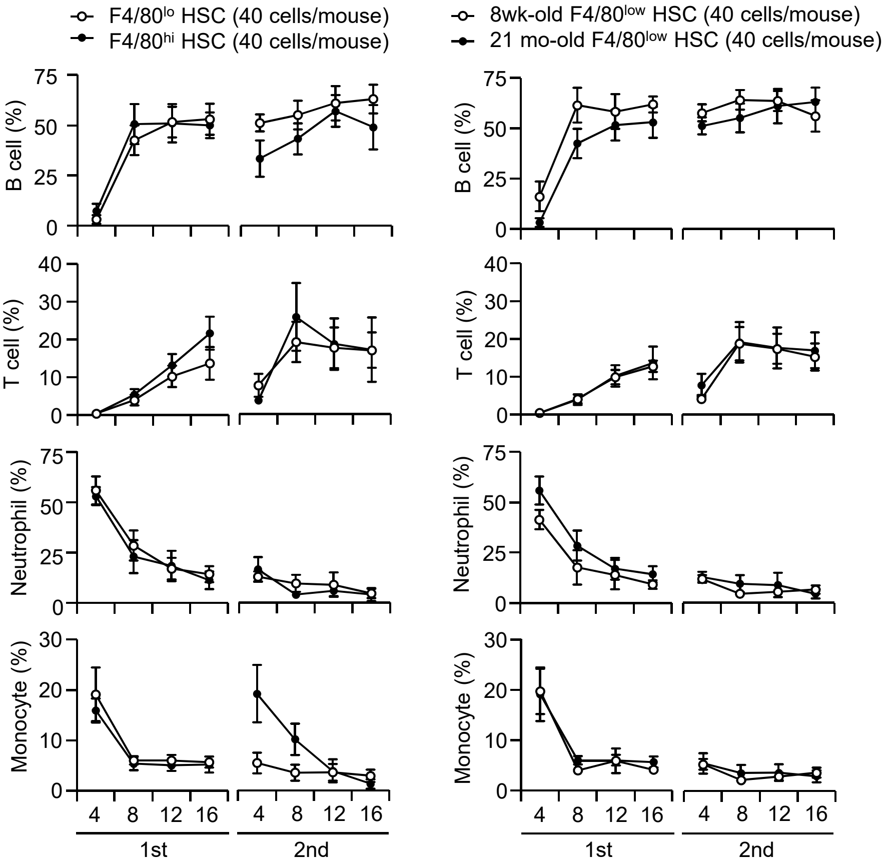


n

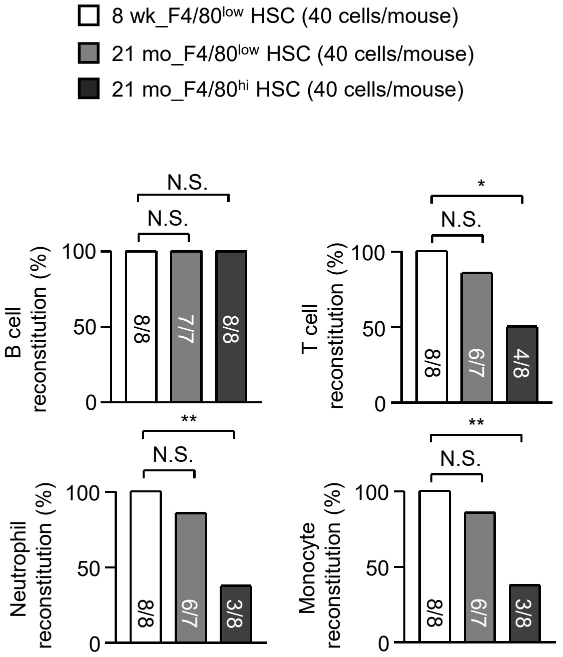

**Supplementary Fig. 7: F4/80$^{lo}$ HSCs during sepsis.**

**a** Cell numbers of CD86$^{-}$ and CD86$^{+}$ HSC in young (8 week-old) and aged (21 month-old) mice. n=13 for 8 week-old and n=9 for 21 month-old mice. **b** Chimerism of donor HSC-derived cells in the blood. CD86$^{-}$ or CD86$^{+}$ HSCs obtained from young (8 week-old) mice or from aged B6J mice (2 year-old) mice were transplanted into lethally irradiated recipient SJL mice. n=8 for 8 week-old and n=7 for 2 year-old mice. **c** Expression of *Actb* and actin rearrangement-associated genes in LT-HSCs obtained from young and from 21 month-old mice. **d** Expression of dormant-associated genes in 8 week-old and in 21 month-old mice. **e-i** CITE-seq analysis of LT-HSCs obtained from WT mice before (purple in e) and 24 hours after LPS-treatment (5 mg/kg) (orange in e). Expression of cell surface F4/80 and CD169 and gene expression levels of these proteins (f), cell surface expression of HSC markers (g) and expression of cell-cycle-associated genes (h) and dormancy-related genes (i) were examined in LT-HSCs. Data for LT-HSCs from naïve WT mice were shared with Fig. 5. **j, k** Expression of cell cycle-associated genes, listed in red in Fig. 5e, and known dormant HSC markers in LT-HSCs during septic (j) and aged (k) conditions. LT-HSCs were obtained 24 hours after the peritoneal administration of LPS (5 mg/kg) (j) or from 13 or 21 month-old mice (k). l Expression of aged HSC markers in 8 week-old and 21-month-old mice. **m, n** F4/80$^{low}$ or F4/80$^{high}$ HSCs obtained from 8 week-old or 21 month-old mice were transplanted into lethally irradiated recipients. Frequencies of B cells, T cells, neutrophils or monocytes in donor HSC-derived cells obtained from peripheral blood of recipient mice (m). The frequency of mice showed the reconstitution of B cell-, T cell-, neutrophil- or monocyte-lineages were examined at 16 weeks after the 2$^{nd}$ transplantation (n). n=8 for F4/80$^{low}$ HSCs obtained from 8 week-old mice and F4/80$^{high}$ HSCs obtained from 21 month-old mice, n=7 for F4/80$^{low}$ HSCs obtained from 21 month-old mice. ***$p<0.001$, N.S. not significant. Student's t-test [a, n] and $\chi^2$ statistics [n] was used. Data are pooled from two [b, m, n] or three [a] independent experiments.

.
